# Bacterial pattern recognition in *C. elegans* by a nuclear hormone receptor

**DOI:** 10.1101/2022.07.12.499718

**Authors:** Nicholas D. Peterson, Samantha Y. Tse, Qiuyu Judy Huang, Celia A. Schiffer, Read Pukkila-Worley

## Abstract

Pattern recognition of bacterial products by host receptors is essential for innate immunity in many metazoans. Curiously, the nematode lineage lost canonical mechanisms of bacterial pattern recognition. Whether other immune receptors evolved in their place is not known. Here, we characterize the first bacterial pattern recognition receptor and its natural ligand in the nematode *Caenorhabditis elegans*. We show that the *C. elegans* nuclear hormone receptor NHR-86/HNF4 senses phenazine-1-carboxamide (PCN), a metabolite produced by pathogenic strains of *Pseudomonas aeruginosa*. PCN binds to the ligand-binding domain of NHR-86/HNF4, a ligand-gated transcription factor, and activates an anti-pathogen transcriptional program in intestinal epithelial cells that provides protection against *P. aeruginosa*. These data de-orphan a nuclear hormone receptor and demonstrate that surveillance of metabolite signals from bacteria allows nematodes to identify virulent pathogens in their environment that are poised to cause disease.

## INTRODUCTION

Many animals rely on pattern recognition to detect microbial infections and activate innate immune defenses (Janeway, 1989). Pattern recognition receptors, such as members of the Toll-like receptor (TLR) protein family, detect physical features of bacterial pathogens, known as pathogen-associated molecular patterns (PAMPs) (Fitzgerald and Kagan, 2020). In mammals, insects, and primitive metazoans such as cnidarians, binding of a PAMP ligand to its host receptor activates protective innate immune defenses that are essential for survival during infection (Brennan and Gilmore, 2018; Brennan et al., 2017). Cnidarians are a common ancestor shared between the nematode *Caenorhabditis elegans* and mammals. Thus, it is surprising that *C. elegans* lacks functional homologs of known bacterial pattern recognition receptors and have lost key intracellular signaling regulators that function downstream of mammalian TLRs, including myeloid differentiation primary-response protein 88 (MYD88), IκB kinase (IKK), and nuclear factor-κB (NF-κB) (Brennan and Gilmore, 2018; Irazoqui et al., 2010). It is unknown whether other immune receptors evolved to replace TLR signaling in nematodes.

*C. elegans* express an expanded family of nuclear hormone receptors compared to other metazoans—274 are present in *C. elegans*, whereas *Drosophila* and humans have only 21 and 48, respectively (Sluder and Maina, 2001; Sluder et al., 1999; Taubert et al., 2011). The marked expansion of this protein family suggests that these proteins have important roles in nematode physiology; however, very few *C. elegans* nuclear hormone receptors have been characterized in detail and the ligands for only four have been determined (Folick et al., 2015; Lin and Wang, 2017; Magner and Antebi, 2008; Motola et al., 2006; Warnhoff et al., 2017; Watson and Walhout, 2014). Interestingly, 259 of the 274 nuclear hormone receptors expanded from a common ancestor shared with the human nuclear hormone receptor, hepatocyte nuclear factor 4 (HNF4) (Sluder and Maina, 2001; Sluder et al., 1999).

Here, we demonstrate that a member of the HNF4 family of nuclear hormone receptors in *C. elegans* is a pattern recognition receptor that directly senses a pathogen-derived metabolite to activate anti-pathogen defenses. Phenazine-1-carboxamide (PCN), a metabolite produced by *Pseudomonas aeruginosa,* binds to the ligand-binding domain of *C. elegans* NHR-86/HNF4 and activates a transcriptional program that provides protection from bacterial killing. These data suggest that the nuclear hormone receptor family may have replaced the function of canonical pattern recognition receptors, which were lost during nematode evolution. In addition, this study reveals that *C. elegans* evolved a mechanism to interpret bacterial metabolites to identify virulent pathogens in its environment that have grown to dangerous levels and are poised to cause disease, a discovery that informs the evolution of innate immune sensing in all metazoans.

## RESULTS

### The pathogen-derived metabolite phenazine-1-carboxamide (PCN) activates anti-pathogen defenses in the *C. elegans* intestine

To determine how *C. elegans* senses infection by the bacterial pathogen *Pseudomonas aeruginosa*, we examined *P. aeruginosa* strains with mutations in key transcriptional regulators that control pathogen virulence (Fig. 1A, Fig. S1A-C). For these studies, a transgenic *C. elegans* strain that carries a GFP-based transcriptional reporter for infection response <u>g</u>ene (*irg*)-4, a secreted immune effector that is transcriptionally induced during bacterial infection, was used as an *in vivo* sensor of immune activation (Anderson et al., 2019; Cheesman et al., 2016; Foster et al., 2020; Peterson et al., 2019; Pukkila-Worley et al., 2012; Pukkila-Worley et al., 2014; Shapira et al., 2006; Troemel et al., 2006). Mutations in three of the 17 *P. aeruginosa* transcriptional regulators eliminated the induction of *C. elegans irg-4*p*::gfp* during infection: pseudomonal mutants in *rhlR*, *pqsR,* and *lasR* (Fig. 1A, Fig. S1A-C). Interestingly, *P. aeruginosa* RhlR, PqsR, and LasR are each transcription factors that function in bacterial quorum-sensing pathways and together control the expression of so-called group behavior genes, which include virulence effectors (Papenfort and Bassler, 2016; Williams and Camara, 2009). Thus, we undertook a secondary screen of 152 *P. aeruginosa* strains with mutations in genes known to be regulated by one of these transcription factors, RhlR (Mukherjee et al., 2017), to identify individual pseudomonal effectors that drive *C. elegans* immune activation (Fig. S1D). We identified only three hits in this screen (*phzA2*, *phzB2*, and *phzH*), all of which contained mutations in phenazine biosynthesis genes (Mavrodi et al., 2001) (Fig. S1E). *C. elegans irg-4*p*::gfp* immune reporter animals infected with a *P. aeruginosa* strain containing clean deletions in both phenazine biosynthesis operons [*P. aeruginosa* Δ*phz* mutant (Dietrich et al., 2006)] fail to upregulate *irg-4*p*::gfp* during infection (Fig. 1A). RNA-sequencing confirmed that *P. aeruginosa* phenazine biosynthesis is required for *C. elegans* innate immune activation (Fig. 1B). Importantly, this experiment identified a group of *C. elegans* genes whose induction during *P. aeruginosa* infection was entirely dependent on the production of phenazines (Fig. 1B). Of these 27 genes, 22 are *C. elegans* innate immune effectors or detoxification genes (Fig. 1B, Table S1A). Examination of transcriptional reporters for the anti-pathogen gene *irg-5* (Fig. S1F) and the cytochrome p450 gene *cyp-35C1* (Fig. S1G) confirmed that the induction of these genes was abrogated during infection with the *P. aeruginosa* Δ*phz* mutant.

**Figure 1.**
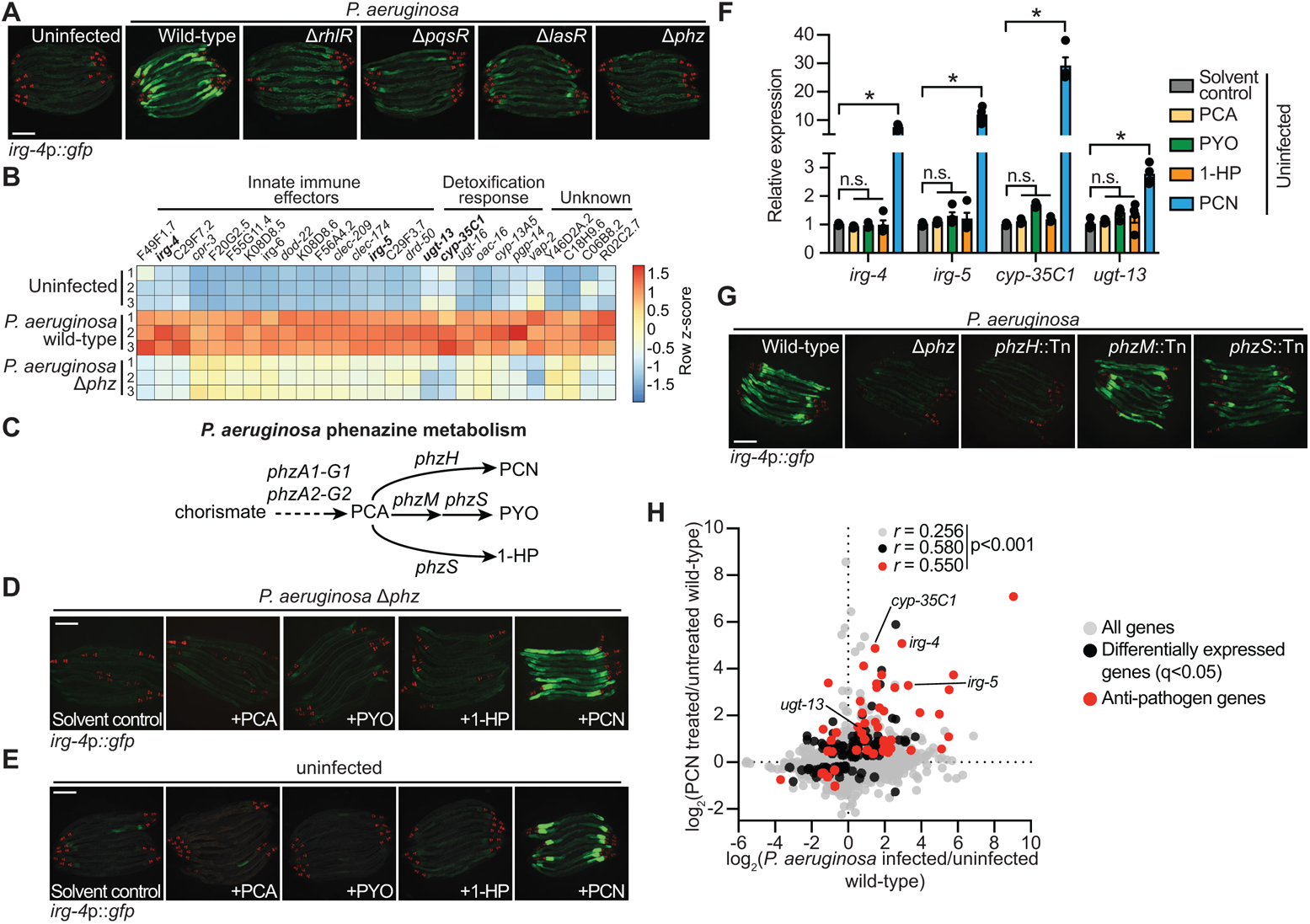
The pathogen-derived metabolite phenazine-1-carboxamide (PCN) activates anti-pathogen defenses in the *C. elegans* intestine. **(A)** Images of *C. elegans irg-4*p::*gfp* transcriptional reporter expression in animals either uninfected or infected with the indicated *P. aeruginosa* strains. **(B)** Heat map of the 27 genes that are induced in *C. elegans* during *P. aeruginosa* infection in a manner dependent on the production of phenazines (q<0.05). Expression level of biological replicates in each condition were scaled by calculating the row z-score for each gene (*n*=3). See also Table S1A. **(C)** A schematic of *P. aeruginosa* phenazine metabolism (PCA, phenazine-1-carboxylic acid; PCN, phenazine-1-carboxamide; PYO, pyocyanin; 1-HP, 1-hydroxyphenazine). **(D and E)** Images of *C. elegans irg-4*p::*gfp* animals during infection with *P. aeruginosa* Δ*phz* **(D)** or grown under standard conditions (uninfected) on media that was supplemented with the indicated phenazines **(E)**. **(F)** qRT-PCR analysis of the indicated anti-pathogen genes in wild-type animals exposed to the indicated phenazines in the absence of infection. Data are the average of biological replicates with error bars giving SEM (*n*=4). *equals p<0.05 (Brown-Forsythe and Welch ANOVA with Dunnett’s T3 multiple comparisons test). Concentration of phenazines used in **D**, **E** and **F** are 112 μM PCN, 112 μM PCA, 119 μM PYO, and 25 μΜ 1-HP. **(G)** Images of *C. elegans irg-4*p::*gfp* animals infected with the indicated *P. aeruginosa* strains. **(H)** Data from mRNA-sequencing experiments comparing genes differentially regulated in wild-type animals exposed to PCN (y-axis) with genes differentially expressed in wild-type animals during *P. aeruginosa* infection (x-axis). All genes are shown in gray. Genes that are differentially expressed in both datasets are shown in black (q<0.05), and the differentially expressed genes annotated as anti-pathogen genes (innate immune effector or detoxification genes) are shown in red. The Pearson correlation coefficient (*r*) between the indicated transcriptional signatures is shown. The location of the representative genes *irg-4*, *irg-5*, *cyp-35C1*, and *ugt-13*, are shown. Scale bars in all images equal 200 μm. See also Table S1B and Fig. S1.

*P. aeruginosa* produces four major phenazine molecules phenazine-1-carboxylic acid (PCA), phenazine-1-carboxamide (PCN), pyocyanin (PYO), and 1-hydroxyphenazine (1-HP) (Fig. 1C) (Mavrodi et al., 2001). Importantly, supplementation with PCN, but not the three other secreted phenazine metabolites, was sufficient to restore both *C. elegans irg-4*p::*gfp* (Fig. 1D) and *cyp-35C1*p::*gfp* (Fig. S1H) activation in the *P. aeruginosa* Δ*phz* mutant. Additionally, in the absence of infection, supplementing PCN, but not the three other phenazines, drove the dose-dependent activation of *C. elegans irg-4*p::*gfp* (Fig. 1E, Fig. S1I) and *cyp-35C1*p::*gfp* expression (Fig. S1J); a finding that was confirmed by qRT-PCR analysis for these and other innate immune effectors (Fig. 1F).

Consistent with the role of PCN in inducing *C. elegans* innate immune defenses, infection with a *P. aeruginosa* strain containing a mutation in *phzH*, a glutamine amidotransferase that synthesizes PCN (Fig. 1C), abrogated the induction of *C. elegans irg-4*p::*gfp* (Fig. 1G), *irg-5*p::*gfp* (Fig. S1K), and *cyp-35C1*p::*gfp* expression (Fig. S1L). We used liquid chromatography-mass spectrometry (LC-MS/MS) to confirm that the *P. aeruginosa phzH* mutant is deficient in the production of PCN but not the other phenazine molecules (Fig. S1M). Notably, *C. elegans* infected with *P. aeruginosa* strains with mutations in either *phzM* or *phzS*, the enzymes that synthesize PYO and 1-HP (Fig. 1C), did not affect the induction of these immune effectors (Fig. 1G, Fig. S1K and L). Moreover, we found that the transcriptional signature of *C. elegans* exposed to PCN mimics that of animals infected with *P. aeruginosa* (Fig. 1H, Table S1B). Thus, the *P. aeruginosa* metabolite PCN specifically and robustly activates *C. elegans* intestinal innate immune defenses.

### The anti-pathogen transcriptional program induced by PCN requires the *C. elegans* nuclear hormone receptor *nhr-86*

We previously demonstrated that the nuclear hormone receptor NHR-86, a ligand-gated transcription factor, surveys the chemical environment to activate intestinal immune defenses (Peterson et al., 2019). To determine if a nuclear hormone receptor senses the pathogen-derived metabolite PCN, we used RNAi and the *C. elegans irg-4*p::*gfp* immune reporter to screen the 274 *C. elegans* nuclear hormone receptor genes (Fig. S2A). Knockdown of only one *nhr*, *nhr-86*, abrogated the induction of *C. elegans irg-4*p::*gfp* (Fig. 2A) and *cyp-35C1*p::*gfp* (Fig. 2B) by PCN treatment and during *P. aeruginosa* infection. Two *nhr-86* loss-of-function alleles *tm2590* (Arda et al., 2010) and *ums12* (Peterson et al., 2019) fully suppressed the induction of *irg-4*p::*gfp* (Fig. 2C) and *irg-5*p::*gfp* (Fig. S2B) under these conditions. We used CRISPR-Cas9 to tag *nhr-86* with an auxin-inducible degron (AID) at its endogenous locus. Treatment with the phytohormone auxin in a transgenic *C. elegans* strain expressing the auxin-binding receptor transport inhibitor response 1 (TIR1) targets NHR-86::AID for degradation by the proteasome (Zhang et al., 2015). We confirmed that auxin treatment induced the degradation of NHR-86::AID protein in this strain (Fig. S2C). Depletion of NHR-86 abrogated the induction of anti-pathogen effector genes following exposure to PCN (Fig. 2D-G) and during *P. aeruginosa* infection (Fig. 2H-K). Consistent with these data, RNA-sequencing revealed that *nhr-86* is required for the induction of *C. elegans* genes following exposure to PCN (Fig. 2L, Fig. S2D, Table S1C). Moreover, of the 133 genes significantly upregulated by PCN in wild-type worms (q<0.05), 63 require *nhr-86* for their induction (Fig. 2L, Fig. S2D, Table S1C).

**Figure 2.**
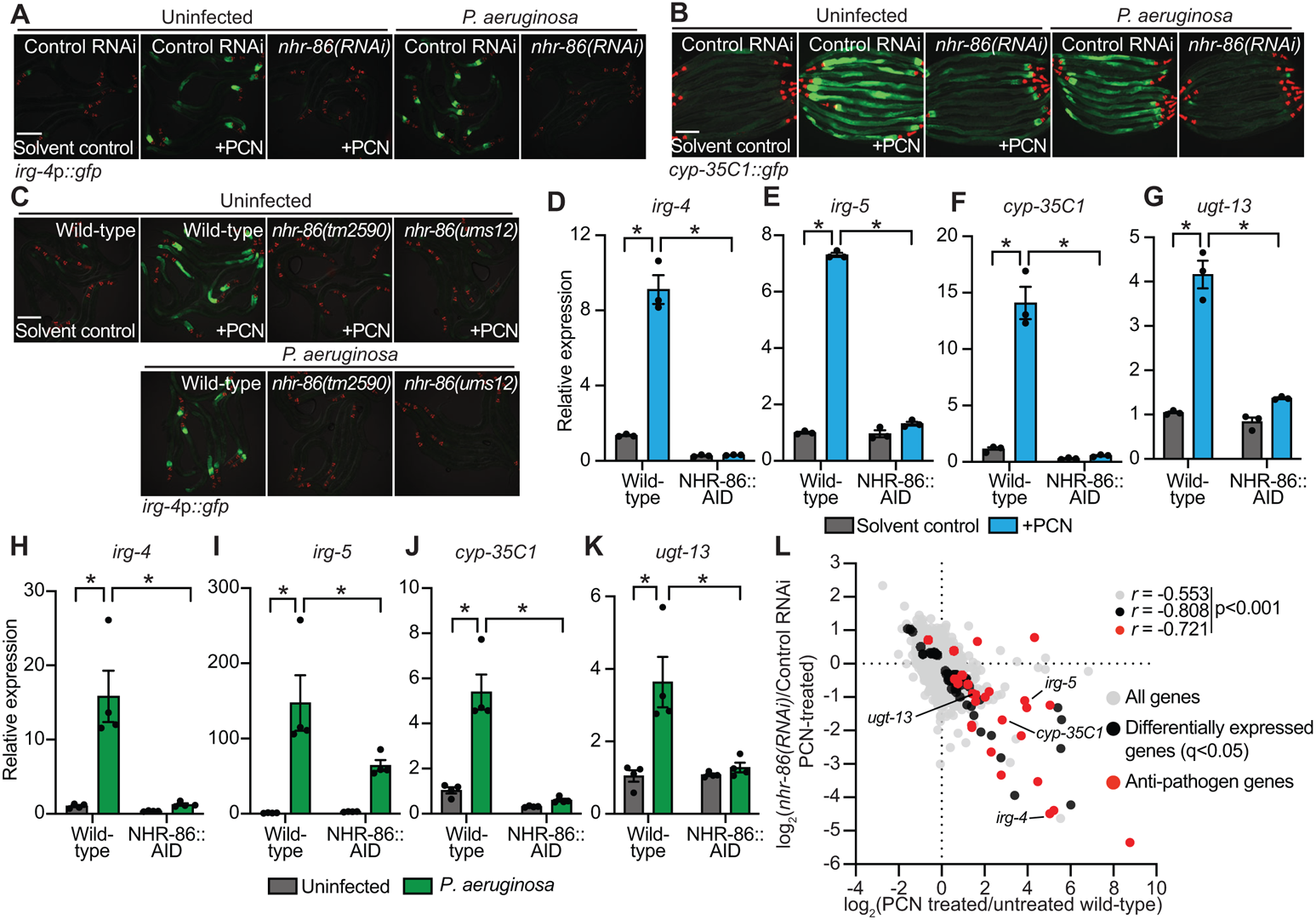
The anti-pathogen transcriptional program induced by PCN requires the *C. elegans* nuclear hormone receptor *nhr-86*. **(A and B)** Images of *C. elegans irg-4*p::*gfp* **(A)** and *cyp-35C1*::*gfp* **(B)** transcriptional reporters with indicated RNAi conditions either exposed to PCN in the absence of infection or during *P. aeruginosa* infection. **(C)** Images of *C. elegans irg-4*p::*gfp* transcriptional reporters with indicated genotypes and conditions. Scale bars in all images equal 200 μm. **(D-K)** qRT-PCR analysis of the indicated innate immune genes in wild-type and NHR-86::AID animals exposed to either PCN in the absence of infection (*n*=3) **(D-G)** or during *P. aeruginosa* infection (*n*=4) **(H-K)**. Data are the mean of biological replicates with error bars giving SEM. *equals p<0.05 (two-way ANOVA with Tukey’s multiple comparisons test) **(L)** Data from mRNA-sequencing experiments comparing genes differentially regulated in *nhr-86(RNAi)* versus control RNAi-treated animals exposed to PCN (y-axis) are compared with genes differentially expressed in wild-type animals exposed to PCN (x-axis). All genes are shown in gray. Genes that are differentially expressed in both datasets are shown in black (q<0.05), and the differentially expressed genes annotated as anti-pathogen genes (innate immune effector or detoxification genes) are shown in red. The location of the representative genes *irg-4*, *irg-5*, *cyp-35C1*, and *ugt-13* are shown. See also Table S1C and Fig. S2.

In a previous study, we showed that NHR-86 binds to the promoters and activates the transcription of intestinal immune defense genes in the presence of a synthetic immunostimulatory molecule (R24) (Peterson et al., 2019). R24 was originally identified in a screen of 37,214 small molecules for those that provided protection from bacterial infection and activated the transcription of the *C. elegans* innate immune reporters *irg-4*p::*gfp* and *irg-5*p::*gfp*, the same genetic tools that uncovered a role for the *P. aeruginosa*-derived metabolite PCN in *C. elegans* immune activation (Fig. 1A, D, E, G, Fig. S1A–E, Fig. S2B and D). Interestingly, PCN and R24 induce transcriptional signatures that are remarkably similar (Fig. S2E, Table S1D). Furthermore, the *nhr-86*-dependent genes that are induced during PCN treatment and those that are upregulated by *nhr-86* following R24 treatment also correlate significantly (Fig. S2F, Table S1E). These data suggest that both the bacterial metabolite PCN and the xenobiotic R24 activate NHR-86 to induce anti-pathogen defenses.

Individual phenazines produced by *P. aeruginosa* activate the mitochondrial unfolded protein response (UPR^mt^) (Deng et al., 2019) and are detected by chemosensory neurons in *C. elegans* (Meisel et al., 2014). Activation of the *C. elegans* UPR^mt^ by phenazines requires the transcription factor ATFS-1 (Haynes et al., 2010). However, knockdown of *atfs-1* by RNAi did not suppress *irg-4*p::*gfp* induction by PCN (Fig. S2G). Additionally, induction of mitochondrial stress by either treatment with mitochondrial poisons (Fig. S2H) or knockdown of a key mitochondrial protease, *spg-7* (Fig. S2I), did not lead to *irg-4*p::*gfp* induction. Likewise, gene set enrichment analysis of genes differentially expressed in wild-type animals following treatment with PCN did not reveal a signature of a mitochondrial stress response induced by either *spg-7(RNAi)* (Fig. S2J) or in the *atfs-1(et18)* gain-of-function mutant (Fig. S2K). In addition, chemosensation of *P. aeruginosa* secondary metabolites, including PCN, induces the transcription of the TGF-β family member *daf-7* in ASJ chemosensory neurons (Meisel et al., 2014). However, the induction of *C. elegans irg-4*p::*gfp* by PCN occurs independently of *daf-7* (Fig. S2L). Collectively, these data demonstrate that the activation of innate immune defenses by PCN occurs through *nhr-86* and not via previously characterized responses to *P. aeruginosa* phenazines.

### The bacterial metabolite PCN and synthetic immunostimulatory molecule R24 bind to the ligand-binding domain of NHR-86

We performed biophysical assays to determine if PCN and R24 are directly sensed by NHR-86. First, we utilized a cellular thermal shift assay (CETSA), a technique based on the principle that the binding of a ligand to its target stabilizes the protein complex against denaturing and aggregation at higher temperatures (Martinez Molina et al., 2013). We found that treatment with PCN and R24 each thermally stabilize NHR-86 (Fig. 3A-C, Fig. S3A). For these studies, we used CRISPR-Cas9 to insert a 3xFLAG tag at the N-terminus of the NHR-86 protein. As a control, we used a strain expressing a transgene that contains a 3xFLAG-labeled NHR-12 protein (Gerstein et al., 2010), which is the closest nematode paralog of NHR-86 (Sural and Hobert, 2021). Using these strains, we probed for either NHR-86 or NHR-12 in whole-cell lysates using an anti-FLAG antibody. PCN and R24 treatments each led to thermal stabilization of NHR-86 over a range of temperatures (Fig. 3A-C, Fig. S3A). We quantified the area under the curve from biological replicates and found that treatment with PCN and R24 each significantly increased the thermal stability of NHR-86 (Fig. 3C, Fig. S3A). Importantly, R24 and PCN failed to thermally stabilize NHR-12 (Fig. 3D and E, Fig. S3B). In addition, the phenazine metabolite PCA, which does not activate host innate immune defenses (Fig. 1D and E, Fig. S1H and J), did not thermally stabilize NHR-86 (Fig. 3A-C, Fig. S3A).

**Figure 3.**
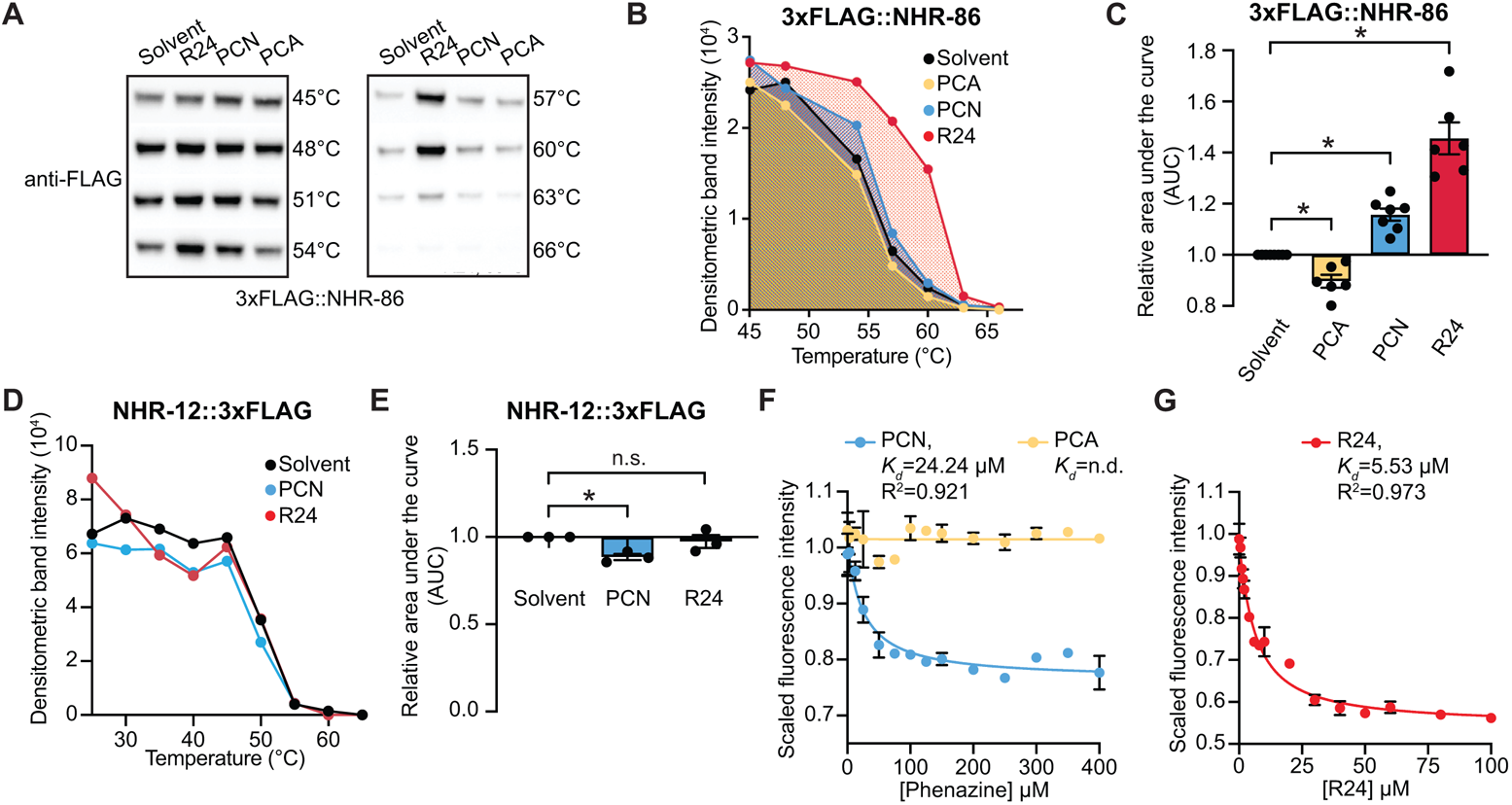
The bacterial metabolite PCN and synthetic immunostimulatory molecule R24 bind to the ligand-binding domain of NHR-86. **(A)** A representative immunoblot of a <u>c</u>ellular thermal <u>s</u>hift <u>a</u>ssay (CETSA) experiment using an anti-FLAG antibody that probed whole cell lysates from a transgenic *C. elegans* strain in which NHR-86 was tagged with 3xFLAG at its endogenous locus. **(B)** A representative densitometric quantification from a CETSA experiment that characterized the interaction of PCN (*n*=7), PCA (*n*=6), and R24 (*n*=6) with 3xFLAG::NHR-86. **(C)** The area under the curve was quantified from each biological replicate experiment in **(B)** and normalized to the solvent control condition. **(D)** Quantification of NHR-12::3xFLAG immunoblot band intensities for each treatment condition and temperature from a representative experiment. **(E)** The area under the curve was quantified from the 3 biological replicate experiments in **(C)** and normalized to the solvent control condition. Data in **C and E** are the average of all biological replicates with error bars giving SEM. *equals p<0.05 (two-tailed, unpaired t-test with Welch’s correction). **(F and G)** Intrinsic tryptophan fluorescence intensity of the purified ligand-binding domain (LBD) of NHR-86 treated with the indicated concentrations of PCN **(F)**, PCA **(F),** and R24 **(G)** each normalized to the solvent control-treated samples. Curves represent a non-linear regression fit of the scaled fluorescence intensity data points for each condition. An equilibrium dissociation constant (*K_d_*) and goodness of fit calculation (R^2^) are shown for each curve. Data in **F and G** are the average of biological replicate samples (*n*=3) with error bars giving SEM. See also Fig. S3.

As an orthologous means to demonstrate that PCN and R24 bind to NHR-86, we expressed and purified the ligand-binding domain (LBD) of NHR-86 from *E. coli* and measured the intrinsic tryptophan fluorescence in the presence of R24 and PCN. Ligand binding to its target protein quenches the fluorescence of the tryptophan residues in the protein (Yammine et al., 2019). PCN (Fig. 3F) and R24 (Fig. 3G) each decreased the intrinsic tryptophan fluorescence intensity of the NHR-86(LBD) in a dose-dependent manner. The equilibrium dissociation constants (*K_d_*) for binding of PCN and R24 to the NHR-86(LBD) are 24.24 μM and 5.53 μM, respectively (Fig. 3F and G). Importantly, PCA, which does not activate host innate immune defenses (Fig. 1D and E, Fig. S1H and J), did not suppress the intrinsic tryptophan fluorescence of the NHR-86(LBD) (Fig. 3F).

To further characterize the binding of R24 and PCN to NHR-86, we modeled the three-dimensional structure of the protein *in silico* (Fig. 4A). HNF4α, the mammalian homolog of *C. elegans* NHR-86, forms a stable homodimer (Chandra et al., 2013), and thus, we used this conformation to model NHR-86. We docked PCN, R24, and PCA into a potential ligand-binding pocket identified in the NHR-86(LBD) (Fig. 4A) and used molecular dynamics simulations to calculate the free energy of binding for these molecules. We found that R24 and PCN each bind stably to NHR-86(LBD), whereas PCA does not (Fig. 4B, Supplemental Videos S1-3). Intriguingly, these calculations also predicted that R24 has an increased affinity for the NHR-86(LBD) compared to PCN, a finding that was confirmed experimentally in both the CETSA thermal stabilization (Fig. 3A-C) and the intrinsic tryptophan fluorescence quenching (Fig. 3F and G) biophysical assays. Consistent with these data, R24 causes more robust induction of anti-pathogen effector genes than PCN at equimolar concentrations (Fig. 4C-F).

**Figure 4.**
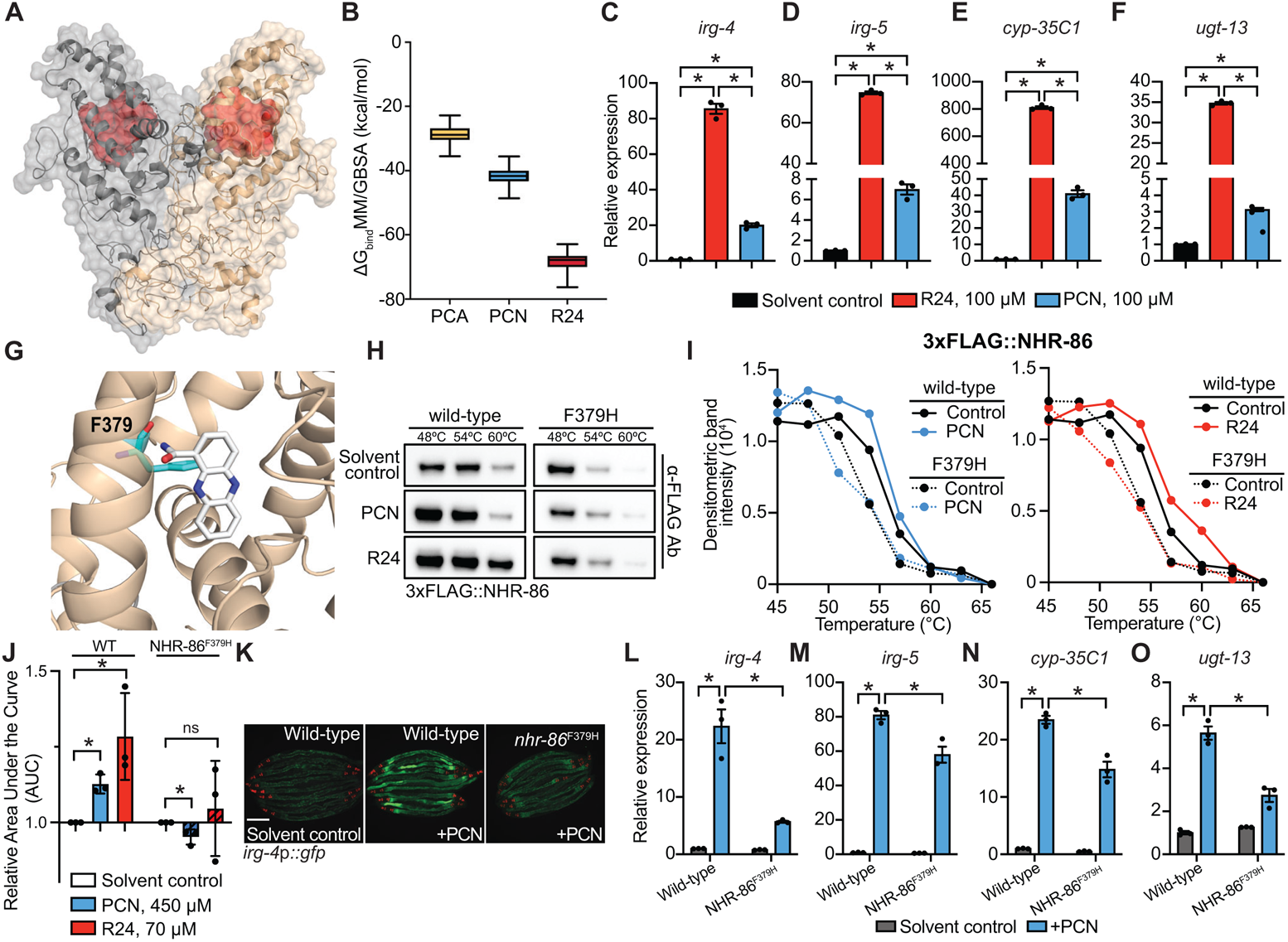
The bacterial metabolite PCN and synthetic immunostimulatory molecule R24 bind to the ligand-binding domain of NHR-86. **(A)** *In silico* molecular modeling of full-length apo NHR-86 as a homodimer. The identified ligand-binding pocket is indicated in red. **(B)** Average free energy of ligand-binding for PCA, PCN, and R24 calculated using the molecular mechanics/generalized Born surface area (MM/GBSA). See also Supplemental Videos 1-3 **(C-F)** qRT-PCR analysis of wild-type animals exposed to either solvent control (1% DMSO) or 100 μM R24 or PCN. Data are the average of biological replicates (*n*=3) with error bars giving SEM. *equals p<0.05 (Brown-Forsythe and Welch ANOVA with Dunnett’s multiple comparisons test). **(G)** An *in silico* model of PCN bound to the identified binding pocket in the NHR-86(LBD). The interaction of phenylalanine 379 (F379) (cyan) and PCN (white) is shown. **(H)** A representative immunoblot of a CETSA experiment using an anti-FLAG antibody that probed whole cell lysates from *C. elegans* 3xFLAG::NHR-86 and 3xFLAG::NHR-86^F379H^ strains treated with indicated conditions. **(I)** A representative densitometric quantification from a CETSA experiment that characterized the interaction of solvent control, PCN, and R24 with 3xFLAG::NHR-86 and 3xFLAG::NHR-86^F379H^ (*n*=3) **(J)** The area under the curve was quantified from each of three biological replicates for the experiment in **(I)** and normalized to the solvent control condition of 3xFLAG::NHR-86. Data are the average of all biological replicates with error bars giving SEM. *equals p<0.05 (two-tailed, unpaired t-test with Welch’s correction). **(K)** Images of indicated *C. elegans irg-4*p::*gfp* animals grown on media that was supplemented with PCN (448 μM). Scale bars in images equal 200 μm. **(L-O)** qRT-PCR analysis of the indicated innate immune genes in wild-type and NHR-86^F379H^ animals exposed to either solvent control or PCN (448 μM) in the absence of infection. Data are the mean of biological replicates (*n*=3) with error bars giving SEM. *equals p<0.05 (two-way ANOVA with Tukey’s multiple comparisons test). See also Fig. S4.

Examination of both PCN and R24 docked *in silico* within the binding pocket of NHR-86(LBD) revealed that the phenylalanine (F) at residue 379 interacts with each of these ligands (Fig. 4G, Fig. S4A). We used CRISPR genome editing to mutate this amino acid (F379H) in *C. elegans* 3xFLAG::NHR-86^F379H^ animals. Importantly, neither PCN nor R24 were able to thermally stabilize 3xFLAG::NHR-86^F379H^ in CETSA experiments performed as described above (Fig. 4H-J, Fig. S4B). Thus, F379 in NHR-86 is required for binding of PCN and R24 to the ligand-binding domain of NHR-86. Consistent with these data, immune effector induction following PCN treatment was attenuated in *C. elegans nhr-86*^F379H^ mutants (Fig. 4K-O). Importantly, we confirmed that 3xFLAG::NHR-86^F379H^ protein was translated at wild-type levels (Fig. S4C).

In summary, these data demonstrate that NHR-86 is a bacterial pattern recognition receptor that directly senses the pathogen-derived metabolite PCN.

### *C. elegans* NHR-86 senses PCN as a marker of pathogen virulence to activate protective anti-pathogen defenses

Phenazine metabolites, in particular PCA and PYO, rapidly kill *C. elegans* in a model of acute pathogen toxicity and are required for the full virulence potential of *P. aeruginosa* in mice (Cezairliyan et al., 2013; Mahajan-Miklos et al., 1999; Recinos et al., 2012). PCN, however, is not toxic to nematodes (Cezairliyan et al., 2013). As previously observed, wild-type *C. elegans* were rapidly killed upon exposure to phenazine toxins secreted into the agar by *P. aeruginosa* (Fig. 5A, Fig. S5A) (Cezairliyan et al., 2013; Mahajan-Miklos et al., 1999). Intriguingly, exposure to the phenazine metabolite PCN provided robust protection from phenazine-mediated killing in this assay (Fig. 5A, Fig. S5B). Post-embryonic degradation of *C. elegans* NHR-86::AID protein abrogated the protection conferred by PCN against toxin-mediated killing (Fig. 5A, Fig. S5B). Thus, sensing of PCN by *C. elegans* NHR-86 activates a protective host response towards secreted phenazine toxins.

**Figure 5.**
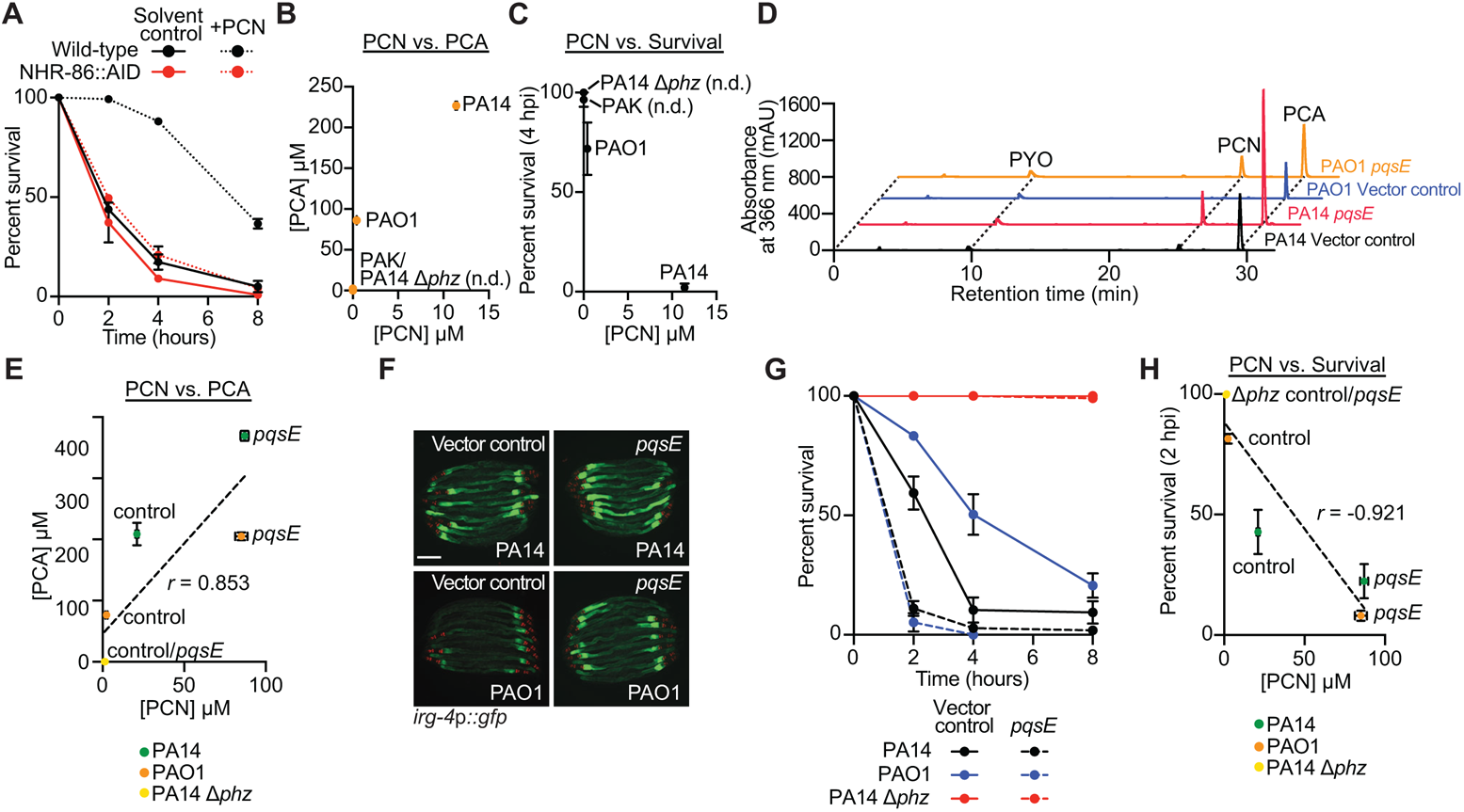
*C. elegans* NHR-86 senses PCN as a marker of pathogen virulence to activate protective anti-pathogen defenses. **(A)** A phenazine toxicity assay in *C. elegans* (also called the “fast kill” assay) with *P. aeruginosa* and *C. elegans* of the indicted genotypes either treated with solvent control or PCN (448 μM). Data are representative of three trials. The difference between PCN-treated wild-type and NHR-86::AID animals is significant (p<0.05, log-rank test, *n*=3). Survival curves for these strains exposed to the *P. aeruginosa* Δ*phz* mutant are shown in Fig. S5A. **(B-D)** HPLC-UV spectroscopy was used to quantify the individual phenazines in the indicated *P. aeruginosa* strains. **(B)** PCN production compared to PCA production in biological replicates of the *P. aeruginosa* strains (*n*=3) **(C)** PCN production is compared to the pathogenicity of *P. aeruginosa* towards *C. elegans* in the phenazine toxicity assay, as quantified by percent nematode survival at four hours. Phenazines not detected (n.d.). **(D)** Liquid chromatography-UV chromatograms of *P. aeruginosa* PA14 or PAO1 strains that express *pqsE* in multicopy (*pqsE*) or a control plasmid (vector control). **(E)** HPLC-UV spectroscopy data showing the comparison of PCN production versus PCA production by biological replicates of the indicated *P. aeruginosa* strains. Pearson correlation coefficient (*r*) is significant (p<0.05, *n*=3). **(F)** Images of *C. elegans irg-4*p::*gfp* animals infected with the indicated *P. aeruginosa* strains. Scale bars in all images equal 200 μm. **(G)** Phenazine toxicity assay with wild-type *C. elegans* and indicated *P. aeruginosa* strains. The difference between the PAO1 control vector and *pqsE* overexpression is significant (p<0.05, log-rank test, *n*=3) **(H)** Comparison of PCN production versus *C. elegans* survival when infected with the indicated *P. aeruginosa* strains in the phenazine toxicity assay. Pearson correlation coefficient (*r*) from biological replicates is significant (p<0.05, *n*=3). Figs. S5B and S5F present the survival data at the four-hour timepoint for the assays in **A** and **G**, respectively. For the assays in **A** and **G**, sample sizes, four-hour survival, and p-values for each replicate are shown in Table S2. See also Fig. S5.

We hypothesized that *C. elegans* utilizes PCN as a marker of pathogen virulence. We found that PCN levels, as quantified by liquid chromatography, in strains of *P. aeruginosa* with varying degrees of virulence potential (PA14, PAO1, and PAK) correlated with the amount of the toxic phenazine PCA produced by these strains (Fig. 5B). Accordingly, the *P. aeruginosa* strains that produced more PCN had enhanced pathogenicity (Fig. 5C) and more robustly induced the *C. elegans* anti-pathogen effectors *irg-4*p::*gfp* (Pukkila-Worley et al., 2014) and *cyp-35*p::*gfp* (Fig. S5C). We drove phenazine production in *P. aeruginosa* PAO1, a strain that naturally produces fewer phenazines (Fig. 5B) and is less pathogenic than PA14 (Fig. 5C), by overexpressing *pqsE*, a pseudomonal gene necessary for phenazine production by the *rhl* quorum-sensing pathway (Garcia-Reyes et al., 2021; Mukherjee et al., 2018; Simanek et al., 2022). Overexpressing *pqsE* in *P. aeruginosa* PAO1, and also in PA14, increased phenazine production (Fig. 5D and E, Fig. S5D and E), including PCN and PCA (Fig. 5D and E, Fig. S5D and E), enhanced the induction of *C. elegans irg-4*p::*gfp* (Fig. 5F), and augmented the pathogenicity of these strains (Fig. 5G and H, Fig. S5F). These data establish a direct connection between phenazine production in *P. aeruginosa,* pathogen virulence potential, and the activation of anti-pathogen defenses in nematodes. Thus, we conclude that *C. elegans* senses the pathogen-derived metabolite PCN to detect toxigenic bacteria in its environment that are poised to cause disease.

## DISCUSSION

Since the recognition that *C. elegans* coordinates inducible immune defenses to provide protection during pathogen infection, the identification of bacterial immune receptors in nematodes has been elusive. Here, we demonstrate that a *C. elegans* nuclear hormone receptor is a *bona fide* pattern recognition receptor that detects a pathogen-derived phenazine metabolite. The phenazine metabolite PCN binds to the ligand-binding domain of NHR-86, which then activates protective anti-pathogen defenses in the intestinal epithelium. We also show that PCN is sensed in *C. elegans* as a specific marker of pathogen virulence, thereby allowing nematodes to interpret secreted bacterial metabolites as a means to identify pathogens in its environment that are poised to cause disease.

It has long been recognized that *C. elegans* senses pathogens indirectly. The G protein coupled receptor DCAR-1 in the *C. elegans* hypodermis recognizes a host ligand, or damage-associated molecular pattern, that is elaborated as a sequela of fungal infection (Zugasti et al., 2014). *C. elegans* also activates immune defenses in response to perturbations in host physiology that accompany infection with pathogenic microbes or the effects of their secreted toxins, a process that is often called surveillance immunity (Dunbar et al., 2012; McEwan et al., 2012; Melo and Ruvkun, 2012; Pellegrino et al., 2014; Pukkila-Worley, 2016). In addition, bloating of the *C. elegans* intestinal lumen induced by microbial colonization activates a behavioral avoidance response and the transcription of immune effector genes (Filipowicz et al., 2021; Singh and Aballay, 2019a, b). Our data now demonstrate *C. elegans* can detect the presence of virulent bacteria specifically through direct detection of a microbial-derived ligand.

Phenazines are produced exclusively by bacteria and play important roles in bacterial physiology (Mavrodi et al., 2006). The production of phenazines is controlled by quorum-sensing pathways (*e.g.*, Las, Rhl, Pqs) that are activated when bacteria reach a high cellular density, such as during biofilm growth (Cezairliyan et al., 2013; Mavrodi et al., 2006; Recinos et al., 2012). In the *P. aeruginosa* biofilm, PCN is a predominant phenazine and functions, in part, as an electron shuttle to promote bacterial respiration within the relatively anoxic environment of the biofilm interior (Recinos et al., 2012; Saunders et al., 2020). These data support our observation that PCN is utilized by *C. elegans* as a specific marker of pseudomonal virulence. Intriguingly, phenazines from *P. aeruginosa* are also sensed to activate innate immunity in mammals. The aryl hydrocarbon receptor AhR recognizes a diverse array of ligands, including environmental toxins, endogenous ligands, and pseudomonal phenazines, to activate protective host defenses (Moura-Alves et al., 2014; Moura-Alves et al., 2019). Thus, the interpretation of bacterial metabolites as a mechanism to direct host defenses towards potential pathogens may be among the most primordial forms of immune sensing in all metazoans.

The *P. aeruginosa* phenazine metabolite PCN activates the transcription of the TGF-β family member *daf-7* in *C. elegans* ASJ chemosensory neurons (Meisel et al., 2014). The activity of *daf-7* in *C. elegans* ASJ neurons is required to program a protective behavioral avoidance response to *P. aeruginosa* (Meisel et al., 2014; Singh and Aballay, 2019a). However, the contribution of PCN in triggering the *C. elegans* behavioral avoidance response is controversial (Meisel et al., 2014; Singh and Aballay, 2019a). Regardless, these data underscore that PCN is a key bacterial molecule that nematodes evolved mechanisms to sense in both neurons and the intestine.

In the absence of infection, the p38 PMK-1 immune pathway in *C. elegans* maintains the constitutive or tonic expression of innate immune defenses (Fletcher et al., 2019; Troemel et al., 2006). Dietary cues, inputs from sensory neurons, and changes in the availability of essential host metabolites, such as cholesterol, adjust the basal activity of the p38 PMK-1 pathway to prime immune effector expression during periods of relative vulnerability to infection (Anderson and Pukkila-Worley, 2020; Cao and Aballay, 2016; Cao et al., 2017; Foster et al., 2020; Peterson et al., 2022; Styer et al., 2008; Wu et al., 2019). During a subsequent bacterial infection, mechanisms of surveillance immunity, intestinal bloating, and NHR-mediated sensing of pathogen-derived metabolites provide nimble mechanisms to direct host defenses specifically toward invading pathogens or secreted toxins (Dunbar et al., 2012; Filipowicz et al., 2021; McEwan et al., 2012; Melo and Ruvkun, 2012; Pellegrino et al., 2014; Peterson et al., 2019; Pukkila-Worley, 2016; Singh and Aballay, 2019a, b).

*C. elegans* consumes bacteria as their primary source of nutrition and lives in microbe-rich habitats like rotting vegetation and compost pits. Thus, direct sensing of structural features of bacteria, such as lipopolysaccharide and flagella, by TLRs may have led to deleterious immune hyperactivation and not provided a selective advantage for nematodes. In this context, we propose that nuclear hormone receptors sense metabolite signals of pathogen virulence, providing an efficient mechanism to identify disease-causing pathogens in the environment. Indeed, a tantalizing hypothesis, which deserves further study, is that the NHR family in *C. elegans* expanded because of their roles in pathogen detection and evolved to replace canonical mechanisms of pattern recognition.

## Supporting information

Movie S1. PCN-NHR-86

Movie S2. R24-NHR-86

Movie S3. PCA-NHR-86

## Acknowledgements

The authors acknowledge Brian Kelch, Ala Shaqra, and Joseph Magrino for reagents and helpful discussions regarding NHR-86 protein expression, Janneke Icso and Paul Thompson for assistance with phenazine quantification, and Melanie Trombly and Merin MacDonald for critical reading of the manuscript. This research was supported by R01 AI130289 (to R.P.W.), R01 AI159159 (to R.P.W.), R21 AI163430 (to R.P.W.), an Innovator Award from the Kenneth Rainin Foundation (to R.P.W.), the Dan and Diane Riccio Fund for Neuroscience (to R.P.W.), F30 AI150127 (to N.D.P.), F30 DK127690 (to S.Y.T.), T32 AI132152 (to N.D.P.), T32 AI095213 (to S.Y.T.), T32 GM107000 (to N.D.P. and S.Y.T.), R01 AI150478 (to C.A.S), R01 GM135919 (to C.A.S.), and R21 AI149716 (to C.A.S.). Some strains were provided by the *Caenorhabditis* Genetics Center, which is funded by the NIH Office of Research Infrastructure Programs (P40 OD010440).

## Author contributions

Conceptualization, NDP, SYT, RPW; Methodology, NDP, SYT, QJY, CAS, RPW; Investigation, NDP, SYT, QJY; Visualization, NDP, SYT, QJY; Funding acquisition, RPW; Project administration, RPW; Supervision, RPW; Writing – original draft, NDP, SYT, RPW; Writing – review & editing, NDP, SYT, QJY, CAS, RPW

## Competing interests

Authors declare that they have no competing interests.

## STAR Methods

### KEY RESOURCES TABLE

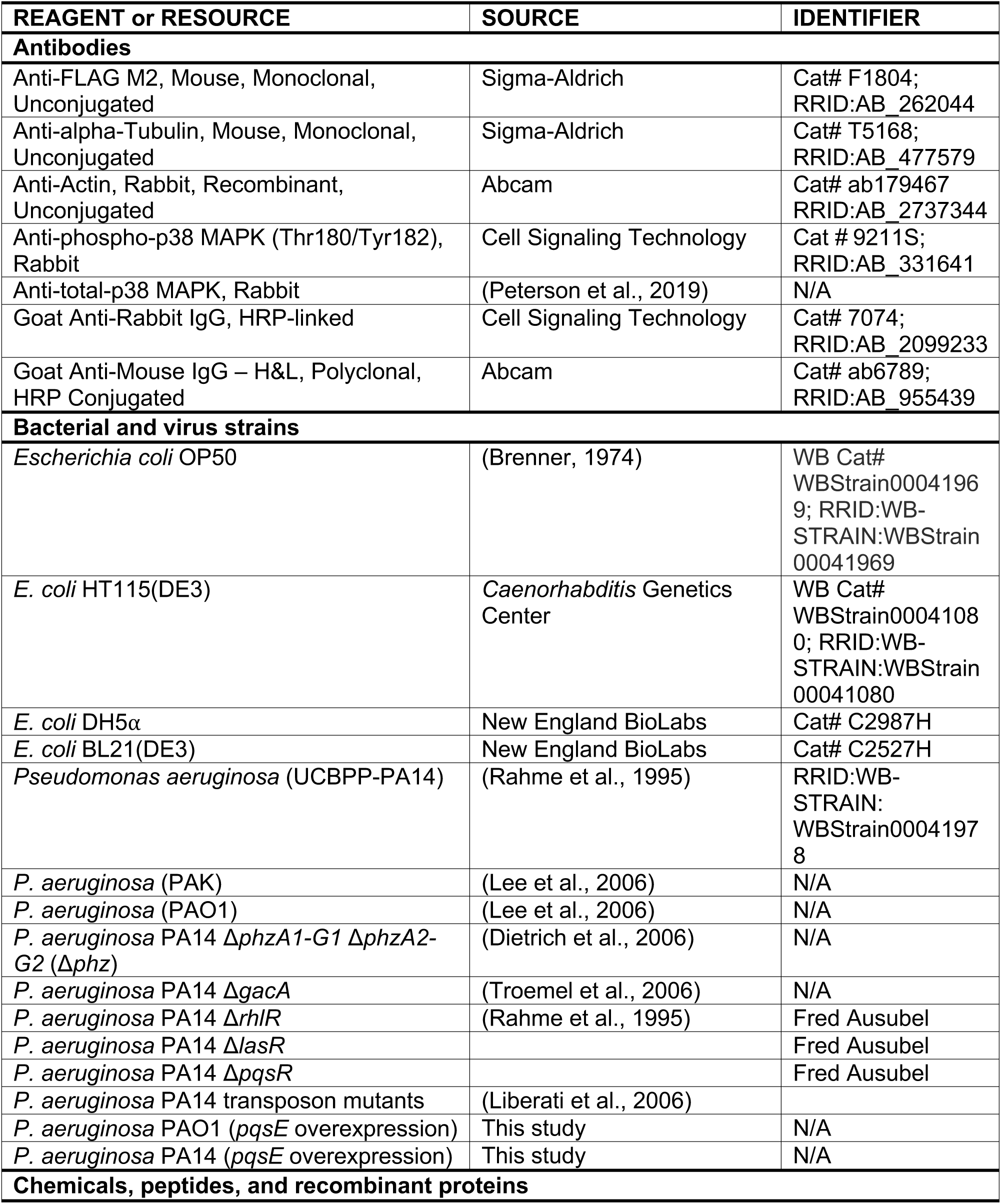
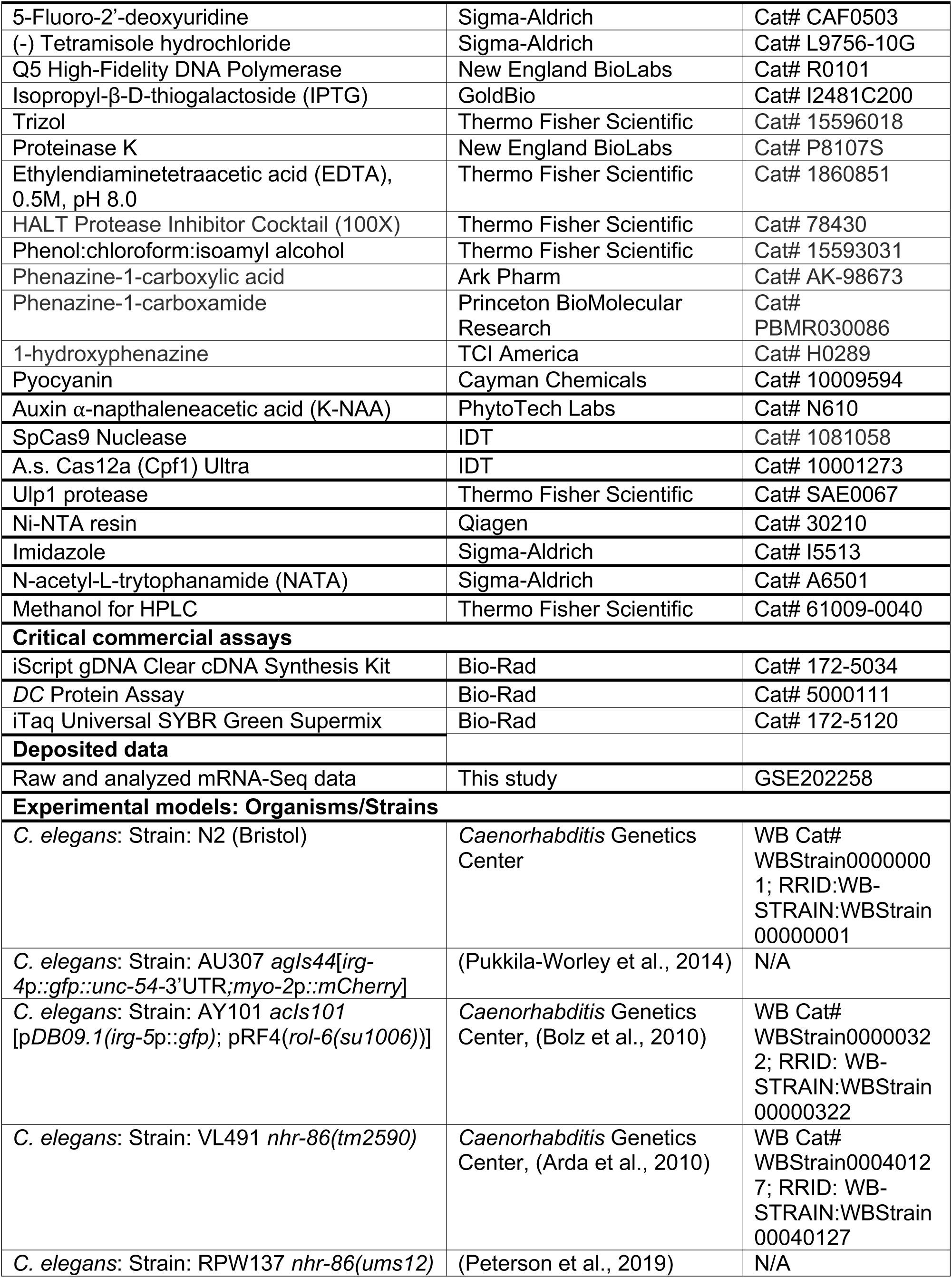
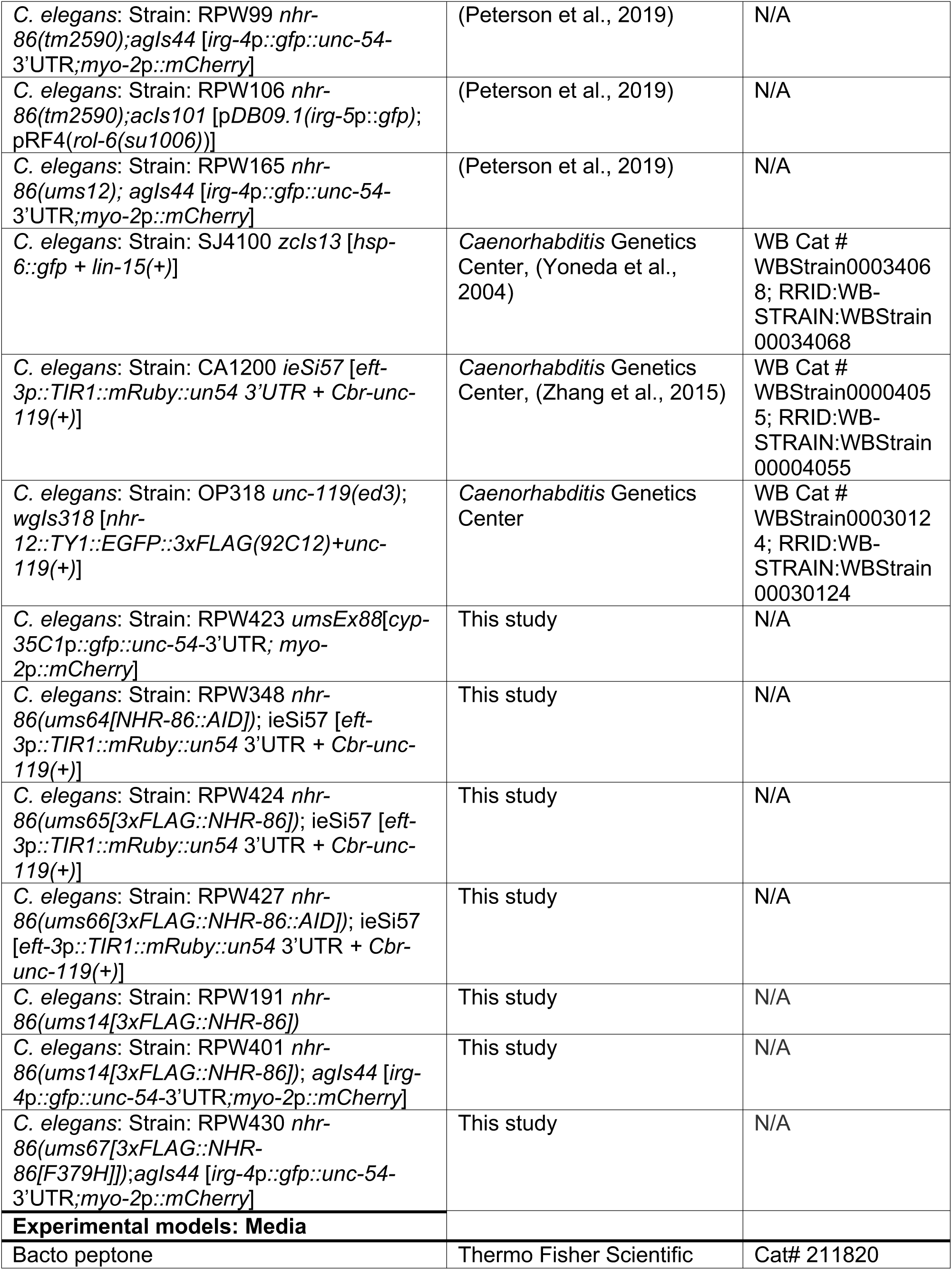
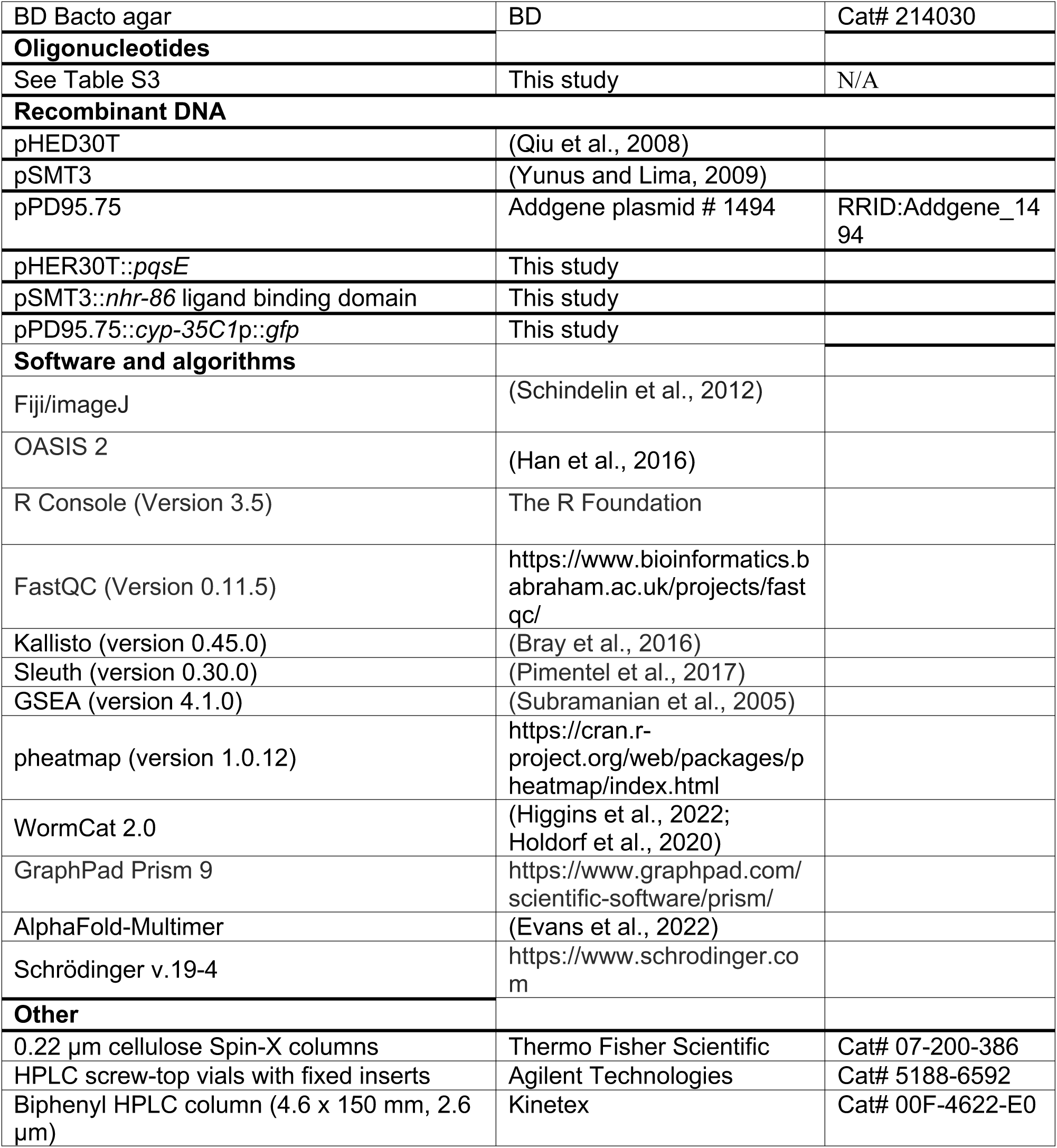

### *C. elegans* and bacterial strains

The previously published *C. elegans* strains used in this study were: N2 Bristol(Brenner, 1974), AU307 *agIs44* [*irg-4*p::*gfp*::*unc-54*-3’UTR*; myo-2*p:*:mCherry*](Pukkila-Worley et al., 2014), AY101 *acIs101* [p*DB09.1(irg-5*p::*gfp)*; pRF4(*rol-6(su1006)*)](Bolz et al., 2010), VL491 *nhr-86(tm2590)*(Arda et al., 2010), RPW137 *nhr-86(ums12)*(Peterson et al., 2019), RPW99 *nhr-86(tm2590)*; *agIs44*(Peterson et al., 2019), RPW106 *nhr-86(tm2590)*; *acIs101*(Peterson et al., 2019), RPW165 *nhr-86(ums12)*; *agIs44*(Peterson et al., 2019), SJ4100 *zcIs13* [*hsp-6::gfp + lin- 15(+)*] (Yoneda et al., 2004), CA1200 *ieSi57* [*eft-3p::TIR1::mRuby::un54 3’UTR + Cbr-unc-119(+)*] (Zhang et al., 2015), OP318 *unc-119(ed3)*; *wgIs318*[*nhr-12::TY1::EGFP::3xFLAG(92C12)+unc-119(+)*] (Gerstein et al., 2010). The strains developed in this study were: RPW423 *umsEx88*[*cyp-35C1p::gfp::unc-54-3’UTR; myo-2p::mCherry*], RPW348 *nhr-86(ums64[NHR-86::AID])*; ieSi57, RPW424 *nhr-86(ums65[3xFLAG::NHR-86])*; ieSi57, RPW427 *nhr-86(ums66[3xFLAG::NHR-86::AID])*; ieSi57, RPW191 *nhr-86(ums14[3xFLAG::NHR-86])*, RPW401 *nhr-86(ums14[3xFLAG::NHR-86])*; *agIs44*, RPW430*nhr-86(ums67[3xFLAG::NHR-86[F379H]])*;*agIs44*. Bacteria used in this study were *Escherichia coli* (*E. coli*) OP50, *E. coli* DH5α, *E. coli* HT115(DE3), and *Pseudomonas aeruginosa* strains PA14 (Rahme et al., 1995), PAO1 (Lee et al., 2006), PAK (Lee et al., 2006), PA14 Δ*phzA1-G1* Δ*phzA2-G2* (Δ*phz*) (Dietrich et al., 2006), PA14 Δ*gacA* (Troemel et al., 2006), and PA14 transposon mutants (Liberati et al., 2006). PA14 Δ*rhlR*, PA14 Δ*lasR* and PA14 Δ*pqsR* were obtained from Fred Ausubel.

#### *C. elegans* growth conditions

*C. elegans* strains were maintained on standard nematode growth medium (NGM) plates [0.25% bacto peptone, 0.3% sodium chloride, 1.7% agar (BD Bacto), 5 μg/mL cholesterol, 25 mM potassium phosphate pH 6.0, 1 mM magnesium sulfate, 1 mM calcium chloride] with *E. coli* OP50 as a food source, as described (Brenner, 1974).

#### Feeding RNAi NHR screen

Knockdown of target genes was performed by feeding *C. elegans E. coli* HT115 expressing dsRNA targeting the gene of interest, as previously described (Conte et al., 2015; Fire et al., 1998; Timmons et al., 2001). In brief, HT115 bacteria expressing dsRNA targeting genes of interest were grown in Lysogeny broth (LB) Lennox medium containing 50 μg/mL ampicillin overnight with shaking (250 rpm) at 37 °C. Overnight cultures were seeded onto NGM containing 5 mM isopropyl β-D-1-thiogalactopyranoside (IPTG) and 50 μg/mL carbenicillin and incubated at 37 °C for 16 hours, after which synchronized L1 animals were transferred to bacterial lawns and allowed to grow until the L4 stage.

For the NHR RNAi screen, *C. elegans irg-4*p*::gfp* transcriptional reporter strains were grown from the L1 to L4 stage on HT115 *E. coli* expressing dsRNA targeting the 274 *C. elegans* NHR genes in the genome. In brief, each well in a 24-well plate containing RNAi agar medium was seeded with 50 µL 5X concentrated overnight culture in M9 buffer of each RNAi clone. Seeded RNAi plates were then incubated overnight at 37 °C. Approximately 50 L1 synchronized *C. elegans irg-4*p::*gfp* transcriptional reporter animals were then dropped onto each bacterial clone and grown until the L4 stage. Animals were then transferred by washing with M9 to 24-well plates containing 25 µg/mL (112 µM) PCN and seeded with 50 µL *E. coli* OP50 for 20 hours before GFP induction was assessed.

#### C. elegans and P. aeruginosa strain construction

##### Strains construction by CRISPR/Cas genome editing

All CRISPR genome editing was performed as previously described (Dokshin et al., 2018; Ghanta and Mello, 2020). CRISPR-Cas9 editing with ssODN homolog directed repair was used to tag *nhr-86* with an auxin inducible degron tag in animals carrying the *ieSi57* transgene, which expresses TIR1 protein in all somatic cells. Animals containing the NHR-86^F379H^ mutation were generated in *nhr-86(ums14[3xFLAG::NHR-86])*;*agIs44* animals using CRISPR-Cas12 directed editing with ssODN homolog directed repair. All CRISPR reagents were purchased from Integrated DNA Technologies. Target guide sequences were selected using the CHOPCHOP web tool (Labun et al., 2019). Single-stranded oligodeoxynucleotide (ssODN) repair templates contained indicated edits, deletions or insertions with 35 bp flanking homology arms. Cas9- and Cas12a- crRNA guide and ssODN sequences are listed in Table S3. The F1 progeny were screened for Rol phenotypes 3 to 4 days after injection and then for indicated edits using PCR and Sanger sequencing. Primer sequences used for genotyping are listed in Table S3.

##### Construction of cyp-35C1p::gfp transgenic reporter animals

Animals carrying the *umsEx88* transgene were constructed as previously described (Pukkila-Worley et al., 2014). Briefly, the region 1000 bp upstream of the *cyp-35C1* 5’UTR was PCR amplified, digested with HindIII and XbaI, and ligated into the *gfp* containing vector pPD95.75. Young adult N2 animals were microinjected with 25 ng/μL *umsEx88* construct along with 5 ng/μL *myo-2*p::*mCherry* co-injection marker. Primer sequences are listed in Table S3.

##### Construction of P. aeruginosa pqsE overexpression strain

*P. aeruginosa pqsE* was amplified by PCR from *P. aeruginosa* PA14 and cloned into the broad host range vector pHERD30T using HiFi DNA Assembly (New England Biolabs). Recombinant plasmids were propagated in *E. coli* DH5α cells and maintained with 50 μg/mL gentamycin selection. *P. aeruginosa* strains were transformed with *pqsE* constructs by electroporation and selected on LB agar containing 50 μg/mL gentamycin, as previously described (Choi et al., 2006). Primer sequences are listed in Table S3.

#### Studies with *C. elegans* GFP-based transcriptional reporters

Immune and detoxification transcriptional reporter assays were performed as previously described (Peterson et al., 2019; Peterson et al., 2022). We previously observed that induction of GFP in the transcriptional reporter *irg-4*p::*gfp* was more robust when the nematode strains were grown on NGM media without supplemented cholesterol (Peterson et al., 2022). Thus, for the studies that utilized *C. elegans irg-4*p::*gfp* animals, NGM was prepared without cholesterol supplementation, and 0.1% ethanol was added to maintain an equivalent ethanol concentration. Single colonies of *P. aeruginosa* strains PA14, PA14 *Δphz*, PA14 transposon mutants, and *pqsE* overexpression strains were grown in 3 mL of LB (for PA14 and PA14 *Δphz*) or LB containing 50 µg/mL gentamicin (for PA14 transposon mutants and *pqsE* overexpression strains) at 37 °C for 14 hours at 250 rpm. 10 µL of culture was then seeded onto “slow-kill” agar (0.35% Bacto-peptone, 0.3% sodium chloride, 1.7% agar, 5 µg/mL cholesterol, 25 mM potassium phosphate, 1 mM magnesium sulfate, 1 mM calcium chloride), allowed to dry, and incubated at 37 °C for 24 hours followed by 25 °C for 24 hours. *E. coli* OP50 was the uninfected control. Phenazines were added to cooled media at the following final concentrations in 1% DMSO, unless otherwise noted: PCA (112 µM, 25 µg/mL), PCN (112 µM, 25 µg/mL), PYO (119 µM, 25 µg/mL), 1-HP (25 µM, 5 µg/mL). Of note, 1-HP was lethal to *C. elegans* when supplemented at a similar concentration as the other phenazines; thus, we performed 1-HP supplementation with the highest concentration that did not affect animal survival in our assay.

*P. aeruginosa Δphz* or *E. coli* OP50 were directly seeded onto phenazine-supplemented plates and dried. For P. aeruginosa *Δphz,* lawns were grown at 37 °C for 24 hours, followed by 25 °C for 24 hours. Around 50–100 *C. elegans* transcriptional reporter animals at the L4 stage were transferred to each bacterial lawn, prepared as described above. Images were taken 20 to 24 hours post-exposure, as described below.

#### Microscopy and image analysis

Nematodes were mounted onto 2% agarose pads, paralyzed with 50 mM tetramisole (Sigma) and imaged using a Zeiss AXIO Imager Z2 microscope with a Zeiss Axiocam 506 mono camera and Zen 2.3 (Zeiss) software. GFP fluorescence in the *irg-4*p::*gfp* transcriptional reporters after infection with *P. aeruginosa* mutants was quantified using the Lionheart FX Automatic Microscope (BioTek Instruments) under a 4X objective. After infection for 24 hours, ∼50 animals were washed three times in M9 buffer containing 0.01% Triton X-100 and transferred to black-sided clear bottom 96-well plates containing 200 µL of 50 mM tetramisole. Animals were allowed to settle for 5 minutes. Individual animals were identified in each well, and mean GFP fluorescence intensity was quantified per animal using the Gen5 software (BioTek Instruments).

#### Gene expression analyses and bioinformatics

RNA-sequencing and data analysis were performed as previously described (Peterson et al., 2022). Briefly, synchronized N2 L1 stage *C. elegans* were grown to the L4 stage on NGM plates seeded with *E. coli* OP50 and transferred by washing with M9 to either *P. aeruginosa, P. aeruginosa Δphz* or *E. coli* OP50 for 4 hours. For the NHR-86 RNA-seq experiment, synchronized L1 stage N2 wild-type animals were grown to L4 on either HT115 L4440 Control RNAi bacteria or HT115 *nhr-86(RNAi)* bacteria. L4 stage animals were then transferred to *E. coli* OP50-seeded agar plates containing solvent control (0.5% DMSO) or 25 µg/mL PCN for 4 hours. For both RNA-seq experiments, animals were harvested by washing with M9, RNA was isolated using TriReagent (Sigma-Aldrich), column purified (Qiagen), and analyzed by 100 bp paired-end mRNA-sequencing using the BGISEQ-500 platform (BGIAmericasCorp) with >20 million reads per sample. The quality of raw sequencing data was evaluated by FastQC (version 0.11.5), clean reads were aligned to the *C. elegans* reference genome (WBcel235) and quantified using Kallisto (version 0.45.0) (Bray et al., 2016). Differentially expressed genes were identified using Sleuth (version 0.30.0) (Pimentel et al., 2017). Pearson correlation statistical analysis was performed using Prism 9.0. Heatmaps of differentially expressed genes were generated using pheatmap (version 1.0.12). Gene set enrichment analysis of RNA-seq was performed using WormCat (Holdorf et al., 2020) for annotation of *C. elegans* gene categories and GSEA (version 4.1.0) (Subramanian et al., 2005) for assessing mitochondrial transcriptional signature in the RNA-seq experiment with PCN.

For the qRT-PCR studies, RNA was reverse transcribed to cDNA using the iScript cDNA Synthesis Kit (Bio-Rad) and analyzed using a CFX384 machine (Bio-Rad) using previously published primers (Cheesman et al., 2016; Peterson et al., 2019; Troemel et al., 2006). All values were normalized against the geometric mean of control genes *snb-1* and *act-3*. Relative expression was calculated using the Pfaffl method (Pfaffl, 2001).

#### Immunoblot analyses

Protein lysates for cellular thermal shift (CETSA) experiments were prepared as described below. For all other immunoblots, protein lysates were prepared using a Teflon Dounce homogenizer from 2,000 *C. elegans* grown to the L4 larval stage on NGM plates seeded with *E. coli* OP50, as previously described (Peterson et al., 2022). LDS Sample Buffer (Thermo Fisher Scientific) was added to a concentration of 1X with 1% β-mercaptoethanol. All samples were incubated at 70 °C for 10 minutes. Total protein from each sample was resolved on NuPage Bis-Tris 4–12% gels (Life Technologies), transferred to 0.2 µM nitrocellulose membranes (Bio-Rad), and blocked with 5% milk in 1x TBS + 0.2% Tween-20 for one hour. Blots were then probed with a 1:1000 dilution of mouse monoclonal anti-FLAG M2 (Sigma, #F1804), mouse monoclonal anti-alpha-Tubulin (Sigma, #T5168), or rabbit monoclonal anti-Actin (Abcam, #ab179467) overnight at 4 °C. Anti-mouse IgG-HRP (Abcam, #ab6789) or anti-rabbit IgG-HRP (Cell Signaling Technology, #7074) secondary antibodies were used at a dilution of 1:10,000 to detect the primary antibodies. Blots were then developed with the addition of SuperSignal™ West Pico PLUS Chemiluminescent Substrate (Thermo Fisher Scientific) and visualized using a ChemiDoc MP Imaging System (Bio-Rad). Band intensities were quantified using ImageJ (Fiji).

#### NHR-86 Ligand-binding domain expression and purification

The NHR-86 ligand-binding domain, codon optimized for *E. coli*, was synthesized by GenScript, amplified by PCR, digested with BamHI and XhoI, and ligated into the vector pSMT3 containing a cleavable N-terminus His6-SUMO tag (Yunus and Lima, 2009). The plasmid was transformed into chemically competent *E. coli* BL21(DE3) cells and maintained with kanamycin selection. For protein expression, a single colony was inoculated into 25 mL LB containing 50 μg/mL kanamycin and grown overnight. Overnight cultures were subcultured to an OD_600_ of 0.05 in Terrific Broth (2.4% yeast extract, 2% bacto tryptone, 0.4% glycerol, 17 mM KH_2_PO_4_, 72 mM K_2_HPO_4_) containing kanamycin and grown at 37 °C with 180 rpm shaking until an OD_600_ of 0.6–0.8. Cells were then placed on ice for 15 minutes. After cooling, isopropyl β-D-1-thiogalactopyranoside (IPTG) was added to a final concentration of 0.5 mM and cultures were incubated for 18 hours at 16 °C with shaking at 180 rpm. Cultures were harvested by centrifugation at 4,000 rpm for 20 minutes at 4 °C, resuspended in binding buffer [50 mM NaH_2_PO_4_ pH 8.0, 500 mM NaCl, 0.001% Tween20, 5 mM β-mercaptoethanol, 10% glycerol (w/v), 5 mM imidazole], flash-frozen in liquid N_2_, and placed at −80°C until purification.

To purify the NHR-86(LBD), samples were thawed and sonicated on ice with a Qsonica Q700 microtip sonicator at an amplitude of 30 for 20 seconds (1 sec on, 1 sec off) followed by 20 seconds off for 12 cycles total. Crude lysate was centrifuged at 10,000 rpm for 30 minutes at 4°C. The soluble fraction was filtered through a 0.45-μM filter and bound to a pre-equilibrated Ni-NTA resin (Qiagen, #30210) by incubating at 4 °C for 1 hour. Bound resin was placed in a column and allowed to flow by gravity. The column was washed with 20 column volumes of wash buffer [50 mM NaH_2_PO_4_ pH 8.0, 500 mM NaCl, 0.001% Tween20, 5 mM β-mercaptoethanol, 10% glycerol (w/v), 20 mM imidazole], and protein was eluted with 5 column volumes of elution buffer [50 mM NaH_2_PO_4_ pH 8.0, 500 mM NaCl, 0.001% Tween20, 5 mM β-mercaptoethanol, 10% glycerol (w/v), 250 mM imidazole]. Protein was dialyzed overnight with 50 mM NaH_2_PO_4_ pH 8.0, 300 mM NaCl, 10% glycerol (w/v). His6-SUMO tag was removed by incubating 7 units of Ulp1 protease (Sigma, #SAE0067) per mg protein with 0.5-mM DTT overnight at 4 °C. Ulp1 protease and His6-SUMO tag were removed by applying protein digestion to a pre-equilibrated Ni-NTA (Qiagen, #30210) column and collecting the flow-through, which was concentrated, dialyzed overnight, flash-frozen in liquid N_2_, and stored at −80 °C.

#### Protein biophysical assays

##### Cellular Thermal Shift Assay (CETSA)

For each assay, approximately 100,000–200,000 L4 3xFLAG::NHR-86 or 3xFLAG::NHR-86^F379H^ animals were resuspended in 1–2 mL of PBS supplemented with HALT protease inhibitor cocktail and lysed using a Teflon Dounce homogenizer on a rotor until all animals were visibly lysed. Cellular debris was removed by centrifugation at 16,000 rpm for 20 minutes at 4°C. Protein in the clarified whole-cell lysate was quantified using the *DC* Protein Assay (Bio-Rad) and adjusted to 10 mg/mL. The whole-cell lysate was then divided into 1.5 mL microcentrifuge tubes and treated with either 1–2% DMSO, 400–500 µM PCA, 400–500 µM PCN, or 70 µM R24 for 15–60 minutes at room temperature. While incubating, 50 µL of lysate from each condition was distributed into PCR tube strips and exposed to increasing temperatures (25–65°C) for 3 minutes on a Bio-Rad C1000 Touch Thermal Cycler, cooled to room temperature for 3 minutes, and immediately placed on ice. Samples were transferred to 1.5-mL microcentrifuge tubes and spun at 20,000 *g* for 20 minutes at 4 °C to remove precipitated proteins. The supernatants for each temperature and condition were carefully transferred to new tubes – without disturbing the pellet or touching the sides of the tubes – containing LDS Sample Buffer (Thermo Fisher) and 1% β-mercaptoethanol. Samples were then assessed for the presence of 3xFLAG::NHR-86 or 3xFLAG::NHR-86^F379H^ using immunoblot analysis, as described above.

##### Intrinsic tryptophan assays

Measurement of NHR-86(LBD) tryptophan fluorescence was performed as previously described with modification (Gao et al., 2018; Yammine et al., 2019). Briefly, 2-μM NHR-86(LBD) protein was incubated with either DMSO (1% final) or increasing concentrations of R24, PCN, or PCA in a 20 μL final volume. Samples were incubated at room temperature for 1 hour in 384 black-walled, round-bottom plates (Corning, #3676). Tryptophan fluorescence was measured using a Molecular Devices SpectraMax iD5 instrument with the following settings: excitation at 295 nm, emission at 340 nm, PMT low, integration 100 ms. To correct for non-specific tryptophan quenching, each compound was simultaneously incubated with 10 μM N-acetyl-L-trytophanamide (NATA) (Sigma, #A6501), a tryptophan analog. The fraction of fluorescence decrease at each compound concentration in NATA was multiplied by the protein solvent control condition, and the measured protein fluorescence at each corresponding compound concentration was then corrected by this factor. Data points for each compound were fit using the following non-linear curve fitting equation:

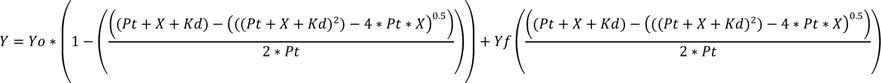

Yo = protein fluorescence intensity with solvent control

Pt = protein concentration

X = concentration of ligand

*K_d_* = equilibrium dissociation constant

#### NHR-86 depletion by auxin-inducible degron

For NHR-86 depletion using the auxin-inducible degron, wild-type and NHR-86::AID animals were treated with 50 µM auxin naphthaleneacetic acid (NAA) (PhytoTech Labs) from the L1 to the L4 larval stage and during all experimental conditions. To avoid NAA impacts on bacterial growth and metabolism, NAA was added on top of bacterial lawns and allowed to diffuse into plates for 2 hours prior to use. We confirmed that NHR-86::AID was degraded during auxin treatment by using CRISPR genome editing to introduce a 3X FLAG tag into the NHR-86::AID strain and immunoblotted for 3xFLAG::NHR-86::AID with an anti-FLAG antibody as described below.

#### *C. elegans* pathogenesis assays

“Fast-killing” *P. aeruginosa* infection experiments were performed as previously described (Cezairliyan et al., 2013; Mahajan-Miklos et al., 1999). In brief, a single colony of *P. aeruginosa* was inoculated into 3 mL of LB Lennox medium and allowed to incubate at 37 °C for 14 hours at 250 rpm. 5 µL of this culture was spread in the center of 35-mm tissue culture plates containing 4 mL of fast-kill agar (1% Bacto-peptone, 1% glucose, 1% sodium chloride, 150 mM sorbitol, 1.7% Bacto-agar). Plates were incubated for 24 hours at 37 °C followed by 24 hours at 25 °C. Approximately 40 L4 larval stage nematodes were transferred to the pseudomonal lawns on fast-kill plates. Dead nematodes were scored at 2, 4, 8 or 24 hours by assessing movement after tapping on the heads with a platinum wire. For the “fast kill” assays with PCN supplementation, agar plates were prepared as above. Following a previously described protocol (Cezairliyan et al., 2013), the bacterial lawn was scraped from the plates after 48 hours of *P. aeruginosa* growth, the agar was melted, and 100 μg/mL PCN or DMSO (1% final) was added to the liquid media. The plates were then re-poured. 20 μL of 20x *E. coli* OP50 was added to plates and allowed to dry. L4 *C. elegans* were then washed with M9 to NGM plates containing 100 μg/mL PCN or DMSO (1% final), prepared as described for studies with transcriptional reporters, for 2 hours before being transferred to supplemented fast-kill plates. Three trials of each pathogenesis assay were performed. Sample sizes, four-hour survival, and p-values for all trials are shown in Table S2.

#### High-Performance Liquid Chromatography-Ultraviolet spectroscopy (HPLC-UV) and Liquid Chromatography/Mass-Spectrometry (LC-MS) quantification of phenazines

Quantification of phenazines was performed as previously described (Schiessl et al., 2019). In brief, agar plates grown with each pseudomonal strain were diced into 5-10 mm cubes and transferred to 50 mL polypropylene tubes containing 5 mL HPLC-grade methanol. Samples were nutated overnight to extract phenazines from both the agar and bacterial biofilms.

Supernatants from methanol-extracted samples were filtered through 0.22 µm cellulose Spin-X columns (Thermo Fisher Scientific 07-200-386), and the filtrates were stored at −80°C until HPLC-UV analysis. On the day of HPLC-UV analysis, 100 µL of supernatant were transferred to HPLC screw-top vials with fixed inserts (Agilent Technologies 5188-6592). Phenazines were quantified using the Agilent 1260 Infinity HPLC with a biphenyl column (Kinetex 00F-4622-E0, 4.6 x 150 mm, 2.6 µm) and a 20 µL injection volume. A gradient method was used, as described previously (Schiessl et al., 2019). Phenazines were quantified by integrating the peaks observed at an absorbance of 366 nm, and phenazines were identified by comparing the retention times to the phenazine standards. All retention times and phenazine quantifications can be found in Table S4.

For figure S2F, *P. aeruginosa* PA14 wild-type and *phzH*::Tn mutants were grown as described above. Bacteria were scraped off the surface of the agar and OD_600_ values were quantified. The agar for each strain was cut up into small pieces and flash-frozen directly in liquid nitrogen. Samples were then pulverized on a Mixer Mill MM 400 (Retsch) under cryogenic conditions at 30 Hz for 90 seconds. Pulverized agar samples were stored at −80 °C until LC-MS/MS analysis.

Phenazines were extracted using methanol and chloroform. The organic phase was dried under nitrogen gas and resuspended in methanol. Samples were filtered through a 0.2 µm PVDF filter and assessed on a Thermo Scientific Ultimate 3000 HPLC system coupled with a Thermo Scientific TSQ Quantiva triple quadrupole mass spectrometer with a Waters Acquity BEH C18 Column and Waters Acquity BEH C18 VanGuard pre-column. The sample injection volume was 2 µL. Mobile phase A consisted of 0.1% formic acid in water, and mobile phase B consisted of 0.1% formic acid in acetonitrile. The gradient started at 35% B for 1.5 min and increased to 99% B over the course of 7 min at a flow rate of 0.25 mL/minute. MS analysis was performed with an electrospray ionization source with a capillary voltage of +3.7 kV. The following was used for the sheath gas: 40 Arb; Aux gas: 10 Arb, vaporizer temperature: 250 °C, ion transfer tube temperature: 325 °C. The multiple reaction monitoring (MRM) parameters were the following: duty cycle time 0.3s, CID gas pressure 1.5 mTorr, Q1 resolution (full width at half maximum, FWHM) 0.7, Q3 resolution (FWHM) 0.7. Quantification of phenazines can be found in Table S4.

#### Molecular modelling and molecular dynamics simulations

AlphaFold-Multimer (Evans et al., 2022) was used to predict the homodimeric structure of full length NHR-86. The model was optimized using Protein Preparation Wizard (Schrödinger v.19-4) to determine protonation states at pH 6.0 and optimize the hydrogen bonding network. A restrained minimization was performed using the OPLS2005 force field (Jorgensen et al., 1996) within an RMSD of 0.3 Å. To determine an optimal binding pocket, SiteMap (Halgren, 2007) (Schrödinger v.19-4) was used with default settings, and the final binding pocket was chosen based on the size, hydrophobicity, and hydrophilicity. Each ligand of interest was converted to an energy minimized 3D molecular structure using LigPrep (Schrödinger v.19-4), and docked within the binding pocket using Glide (Friesner et al., 2004) (Schrödinger v.19-4). Energy minimization was conducted for ligand poses with highest docking scores. A multistage 100 ns molecular dynamics simulation with randomized starting velocities was performed for each ligand-protein complex using Desmond (Schrödinger v.19-4). Forcefield parameters were assigned using OPLS3 (Harder et al., 2016). Each complex was solvated in a cubic box with at least 15 Å between any solute atom and the periodic boundaries using the TIP3P water model. Charges were neutralized using sodium and chloride ions, and additional counterions were added up to concentration of 0.15 M. MM/GBSA calculations were carried out using 100 frames of the simulation using a custom script. Structural figures and movies were generated using PyMOL (v. 2.3.4) and VMD (v. 1.9.4).

#### Quantification and statistical analysis

Differences in the survival of *C. elegans* in the *P. aeruginosa* pathogenesis assays were determined with the log-rank test after survival curves were estimated for each group with the Kaplan-Meier method. OASIS 2 was used for these statistical analyses (Han et al., 2016). Statistical hypothesis testing was performed with Prism 9 (GraphPad Software) using methods indicated in the figure legends. Sample sizes, four-hour survival, and p-values for all trials are shown in Table S2.

#### Data and code availability

The mRNA-seq datasets are available from the NCBI Gene Expression Omnibus using the accession number GSE202258. All other data are available in the manuscript and the accompanying Table S5, which contains all source data and statistical tests used.

**Table S1A-E. (A)** *C. elegans* genes significantly differentially induced during *P. aeruginosa* infection in a phenazine-dependent manner in the RNA-seq experiments presented in Fig. 1B. **(B)** *C. elegans* genes significantly differentially induced by PCN exposure in the absence of infection and during *P. aeruginosa* infection in the RNA-seq experiments presented in Fig. 1H. **(C)** *C. elegans* genes significantly differentially induced by PCN in an *nhr-86*-dependent manner in the RNA-seq experiments presented in Fig. 2L and Fig. S2D. **(D)** *C. elegans* genes significantly differentially induced by PCN and R24 exposure in the absence of infection in the RNA-seq experiments presented in Fig. S2E. **(E)** *C. elegans* genes significantly differentially induced by PCN and R24 in an *nhr-86*-dependent manner in the RNA-seq experiments presented in Fig. S2F.

**Table S2.** Sample sizes, four-hour survival, and p values for the C. elegans pathogenesis assays.

**Table S3.** Primer, crRNA guide and ssODN sequences designed for this study.

**Table S4.** HPLC-UV and LC-MS/MS Phenazine quantification retention times and abundance.

**Table S5.** Source data and statistical tests used for each figure and supplemental figure.

**Video S1.** Molecular dynamics simulation of PCN docked into the predicted ligand-binding pocket of NHR-86(LBD). This simulation was used to calculate the free energy of binding for this molecule, the data for which is shown in Fig. 4B.

**Video S2.** Molecular dynamics simulation of R24 docked into the predicted ligand-binding pocket of NHR-86(LBD). This simulation was used to calculate the free energy of binding for this molecule, the data for which is shown in Fig. 4B.

**Video S3.** Molecular dynamics simulation of PCA docked into the predicted ligand-binding pocket of NHR-86(LBD). This simulation was used to calculate the free energy of binding for this molecule, the data for which is shown in Fig. 4B.

---

**Figure S1.**
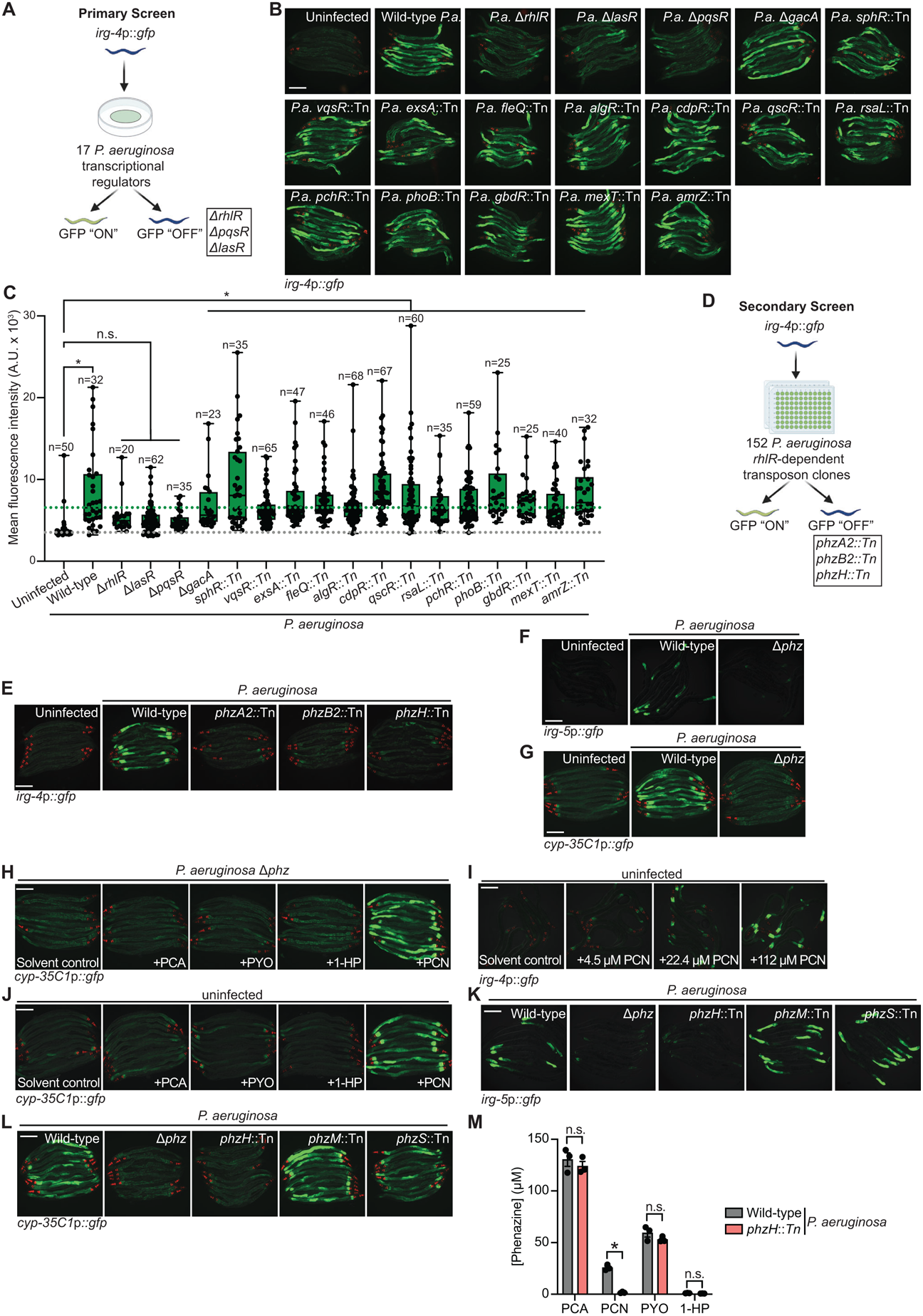
The pathogen-derived metabolite phenazine-1-carboxamide (PCN) activates anti-pathogen defenses in the *C. elegans* intestine. **(A)** A schematic of the primary screen of 17 *P. aeruginosa* virulence-related transcription factors using *C. elegans irg-4*p::*gfp* transcriptional immune reporter animals. Hits are indicated: *rhlR*, *pqsR*, and *lasR*. **(B)** Representative images of *C. elegans irg-4*p::*gfp* transcriptional reporter animals either uninfected or infected with the indicated *P. aeruginosa* strains. **(C)** Quantification of *irg-4*p::*gfp* transcriptional reporter intensity in animals either uninfected or infected with the indicated *P. aeruginosa* strains. Sample size (n) for each strain is indicated. Each data point indicates one animal. Box and whisker plots represent the median with minimum, second quartile, third quartile, and maximum indicated for each condition. *equals p<0.05 (one-way ANOVA with Dunnett’s multiple comparisons test). **(D)** A schematic of the secondary screen of 152 *P. aeruginosa* mutant strains, each with a mutation in a gene dependent on the transcription factor *rhlR* for full expression. Hits are indicated: *phzA2*, *phzB2*, and *phzH*. **(E-G)** Images of the *C. elegans* transcriptional reporters *irg-4*p::*gfp* **(E)**, *irg-5*p::*gfp* **(F)**, and *cyp-35C1*p::*gfp* **(G)** animals either uninfected or infected with the indicated *P. aeruginosa* strains. **(H-L)** Images of *cyp-35C1*p::*gfp*, *irg-4*p::*gfp*, and *irg-5*p::*gfp* transcriptional reporter expression in animals with indicated conditions and as described in Figure 1. **(M)** Quantification of phenazines in *P. aeruginosa* wild-type and *phzH::Tn* replicates by LC-MS/MS. Data are the mean with errors bars giving SEM of biological replicates (*n*=3). *equals p<0.05 (two-way ANOVA with Šídák’s multiple comparisons test). Scale bars in all images equal 200 μm. See also Fig. 1.

**Figure S2.**
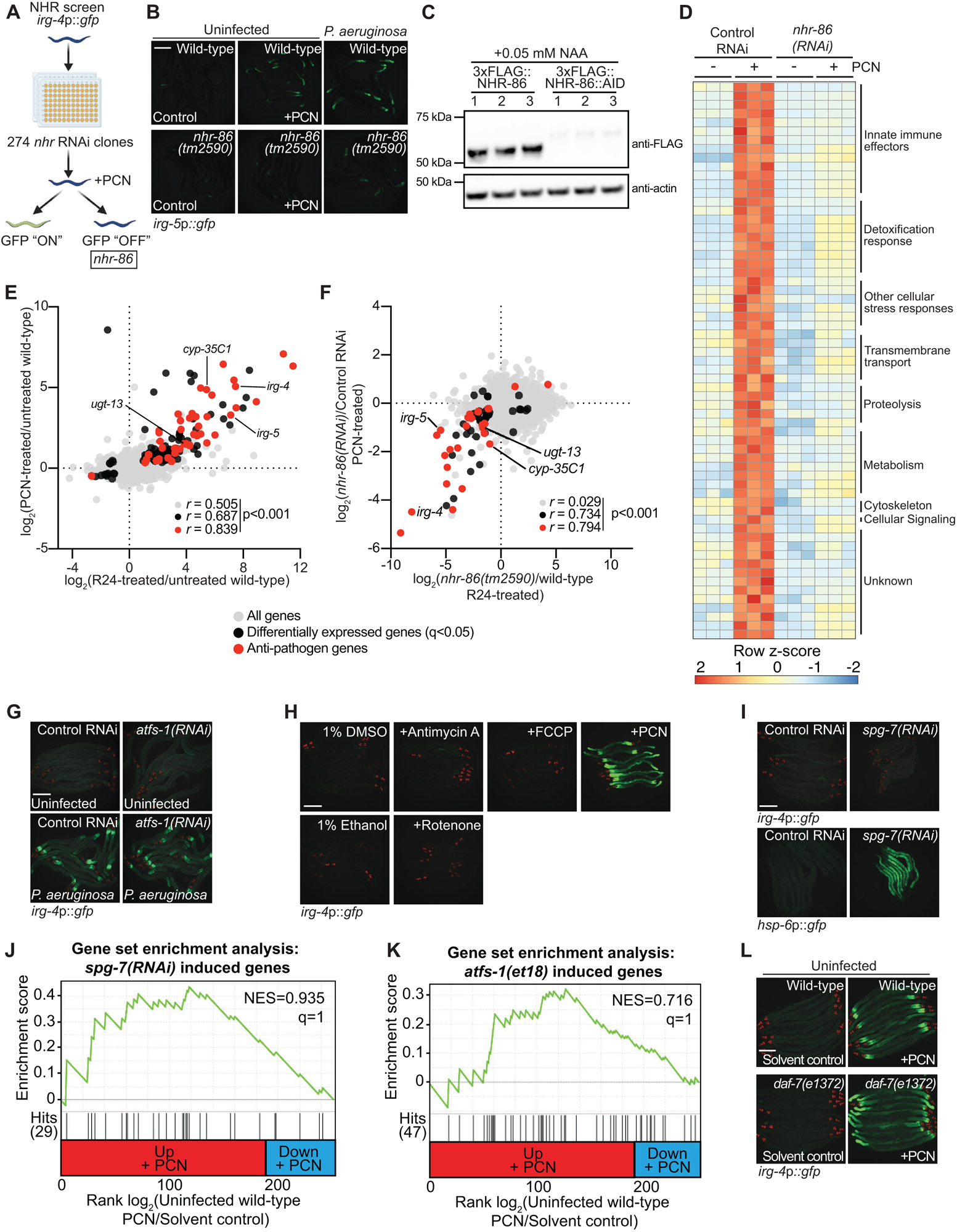
The anti-pathogen transcriptional program induced by PCN requires the *C. elegans* nuclear hormone receptor *nhr-86.* **(A)** A schematic of the screen of 274 *nhr* RNAi clones using *C. elegans irg-4*p::*gfp* transcriptional immune reporter animals. The lone hit is indicated: *nhr-86*. **(B)** Images of *irg-5*p::*gfp* transcriptional reporter expression with indicated genotypes and conditions, as described in Fig. 2C. Scale bars in all images equal 200 μm. **(C)** Immunoblot analysis of lysates from biological replicates of 3xFLAG::NHR-86 and 3xFLAG::NHR-86::AID animals expressing the TIR1 transgene in all somatic tissues treated with 50 μM NAA (*n*=3). Lysates were probed with anti-FLAG and anti-Actin antibodies. The expected size of 3xFLAG::NHR-86 and 3xFLAG::NHR-86::AID is 49.8 kDa and 54.6 kDa respectively. **(D)** A heat map of the 63 genes that are induced in *C. elegans* during PCN exposure in an *nhr-86* dependent manner (q<0.05 RNA-seq analysis, see Materials and Methods). Gene expression in each condition was scaled by calculating a row z-score for each gene. See also Table S1C. **(E)** An mRNA-seq experiment as described in Fig. 1H, except the genes differentially expressed in wild-type animals exposed to R24 in the absence of infection is compared to the genes induced by PCN. See also Table S1D. **(F)** Data from an mRNA-seq experiment as described in Fig. 2L, except the genes that require *nhr-86* for their expression during PCN and R24 treatment are compared. See also Table S1E. **(G)** Images of *C. elegans irg-4*p::*gfp* animals exposed to the indicated RNAi clones and conditions. **(H)** Images of *C. elegans irg-4*p::*gfp* animals exposed to either the solvent controls (1% DMSO or 1% ethanol), the indicated mitochondrial toxins, or PCN. **(I)** Images of *C. elegans irg-4*p::*gfp* or *hsp-6*p::*gfp* reporters treated with *spg-7(RNAi)*. **(J and K)** Gene set enrichment analysis (GSEA) of genes induced in *spg-7(RNAi)* treated animals **(J)** or in the *atfs-1(et18)* gain-of-function allele **(K)** in the RNA-seq of wild-type *C. elegans* exposed to PCN. In **J** and **K**, fold change in the expression of the significantly differentially expressed genes (q<0.05) in uninfected animals exposed to PCN in the absence of infection are ranked from higher expression (red) to lower expression (blue). Normalized enrichment score (NES) and q-value are indicated. Genes induced by either condition and found in the PCN transcriptional profile are indicated by hit number in the left margin and black lines. **(L)** Images of *C. elegans irg-4*p::*gfp* reporter expression with indicated genotypes and conditions. Scale bars in all images equal 200 μm. See also Fig. 2.

**Figure S3.**
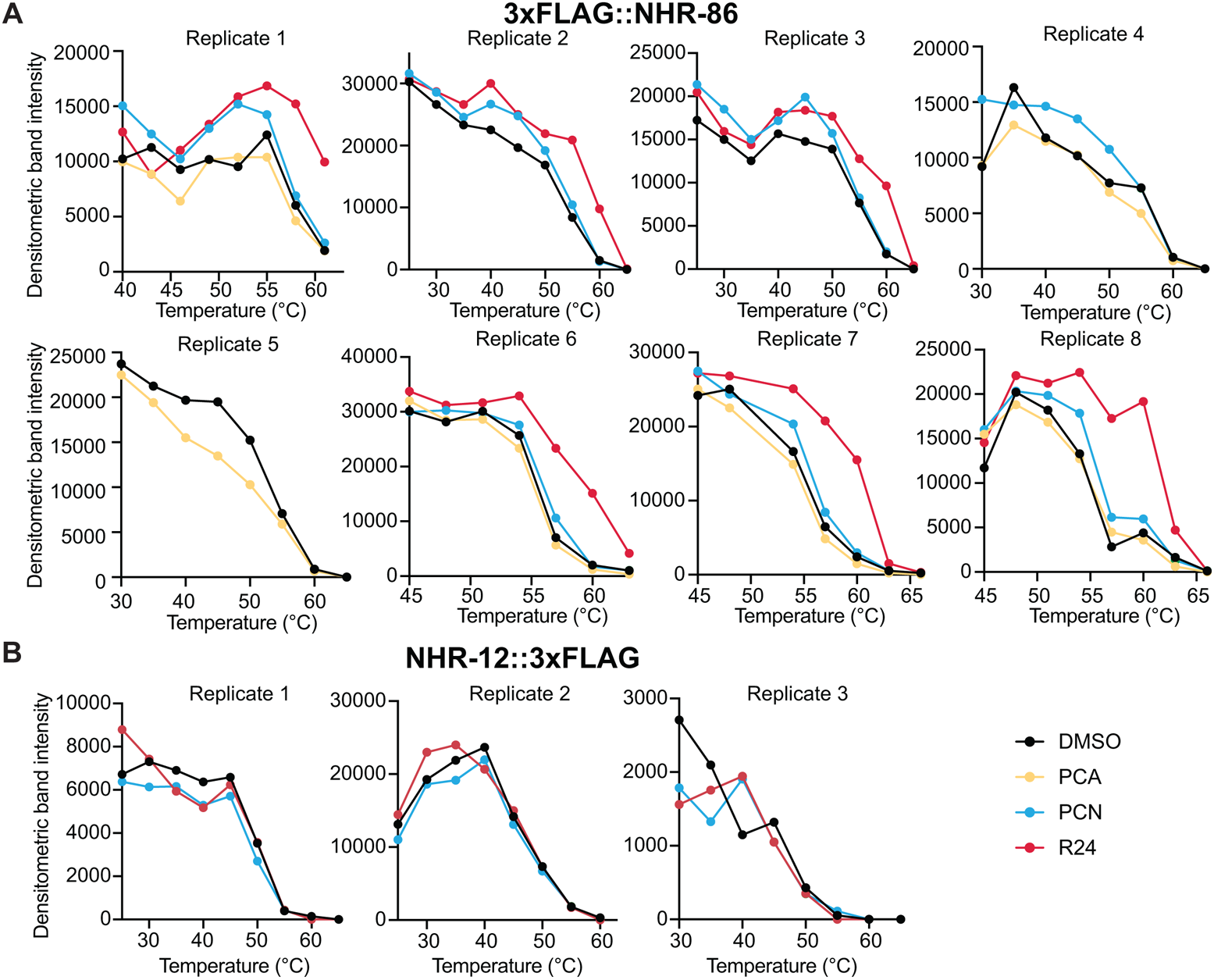
The bacterial metabolite PCN and synthetic immunostimulatory molecule R24 bind to the ligand-binding domain of NHR-86. **(A)** Quantification of 3xFLAG::NHR-86 immunoblot band intensities for each treatment condition and temperature for all replicates. **(B)** Quantification of NHR-12::3xFLAG immunoblot band intensities for each treatment condition and temperature for each replicate. See also Fig. 3.

**Figure S4.**
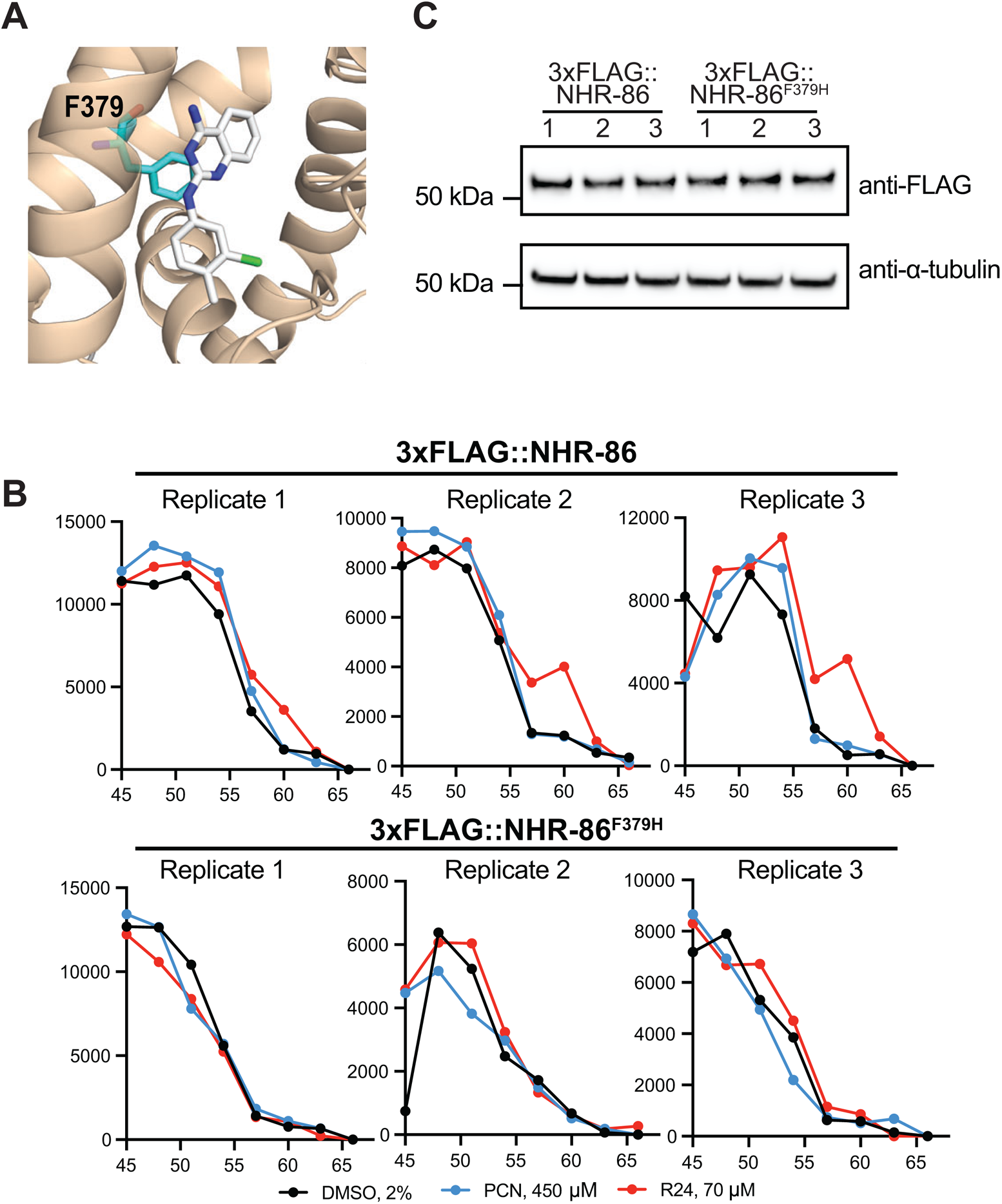
The bacterial metabolite PCN and synthetic immunostimulatory molecule R24 bind to the ligand-binding domain of NHR-86. **(A)** An *in silico* model of R24 bound to the identified binding pocket in the NHR-86(LBD). The interaction of phenylalanine 379 (F379) (cyan) and R24 (white) is shown. **(B)** Quantification of 3xFLAG::NHR-86 and 3xFLAG::NHR-86^F379H^ immunoblot band intensities for each treatment condition and temperature for all replicates. **(C)** Immunoblot analysis of lysates from biological replicates of 3xFLAG::NHR-86 and 3xFLAG::NHR-86^F379H^ animals (*n*=3). Lysates were probed with anti-FLAG and anti-α-tubulin antibodies. The expected size of 3xFLAG::NHR-86 is 49.8 kDa. See also Fig. 4.

**Figure S5.**
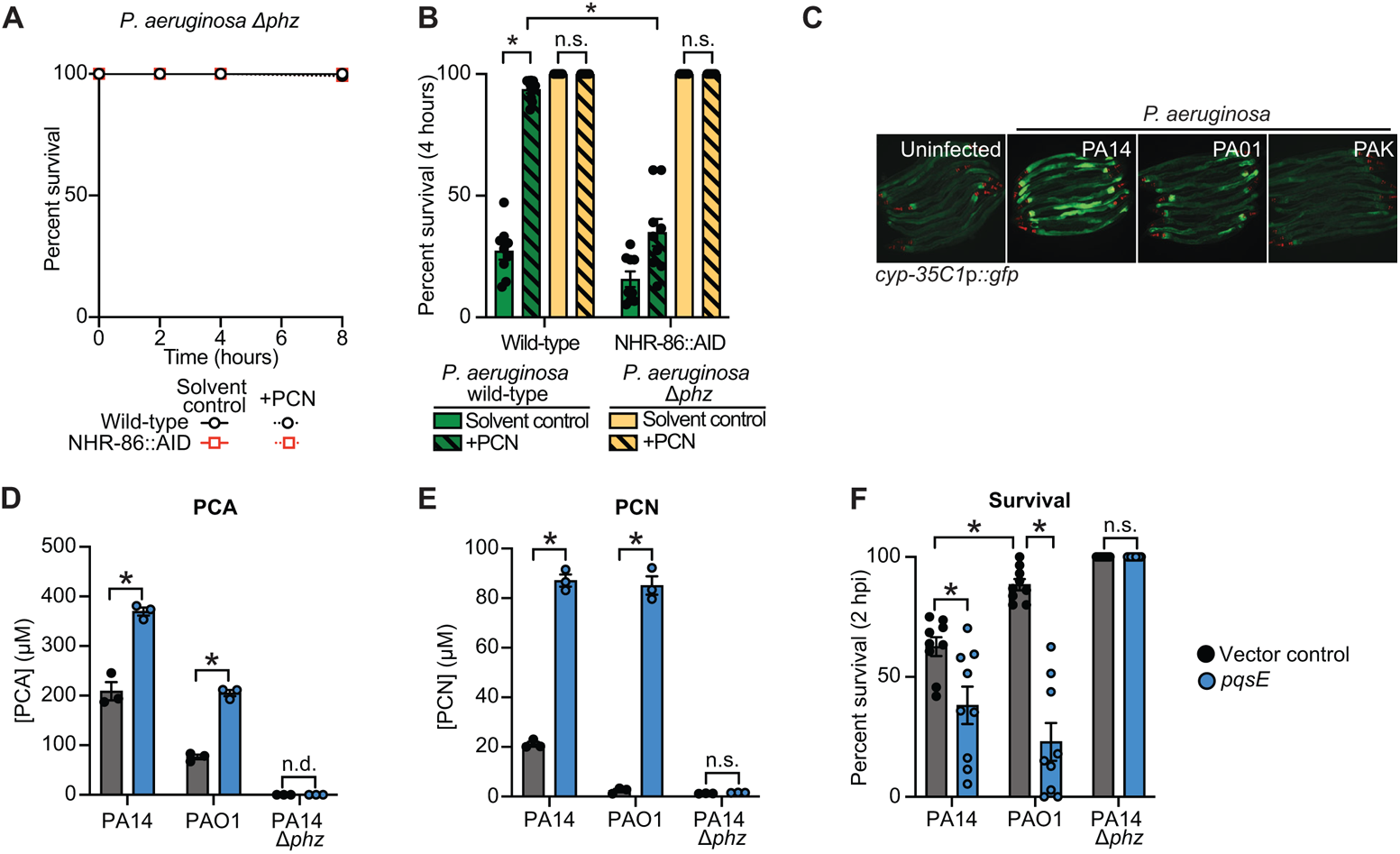
*C. elegans* NHR-86 senses PCN as a marker of pathogen virulence to activate protective anti-pathogen defenses. **(A)** *C. elegans* phenazine toxicity assay with *P. aeruginosa* and *C. elegans* of the indicted genotypes either treated with solvent control or PCN. Data are representative of three trials. Differences between all conditions is not significant (p<0.05). **(B)** Survival of indicated *C. elegans* genotypes four hours after exposure to the indicated conditions. *equals p<0.05 (two-way ANOVA with Tukey’s multiple comparisons test). **(C**) Images of *C. elegans cyp-35C1*p::*gfp* animals either uninfected or infected with the indicated *P. aeruginosa* genotypes. Scale bars in all images equal 200 μm. **(D and E).** Quantification of phenazines PCA **(D)** and PCN **(E)** in indicated *P. aeruginosa* strains by HPLC-UV spectroscopy from biological replicates (*n*=3). Phenazines not detected (n.d.). *equals p<0.05 (two-way ANOVA with Tukey’s multiple comparisons test). **(F)** Survival of wild-type *C. elegans* two hours post-infection with the indicated *P. aeruginosa* genotypes. *equals p<0.05 (two-way ANOVA with Tukey’s multiple comparisons test). Sample sizes, four-hour survival, and p-values for acute phenazine toxicity assays are shown in Table S2. See also Fig. 5.

## Notes

### Competing Interest Statement

The authors have declared no competing interest.

## References

Anderson, S.M., Cheesman, H.K., Peterson, N.D., Salisbury, J.E., Soukas, A.A., and Pukkila-Worley, R. (2019). The fatty acid oleate is required for innate immune activation and pathogen defense in *Caenorhabditis elegans*. PLoS Pathog 15, e1007893.

Anderson, S.M., and Pukkila-Worley, R. (2020). Immunometabolism in *Caenorhabditis elegans*. PLoS Pathog 16, e1008897.

Arda, H.E., Taubert, S., MacNeil, L.T., Conine, C.C., Tsuda, B., Van Gilst, M., Sequerra, R., Doucette-Stamm, L., Yamamoto, K.R., and Walhout, A.J. (2010). Functional modularity of nuclear hormone receptors in a *Caenorhabditis elegans* metabolic gene regulatory network. Mol Syst Biol 6, 367.

Bolz, D.D., Tenor, J.L., and Aballay, A. (2010). A conserved PMK-1/p38 MAPK is required in *Caenorhabditis elegans* tissue-specific immune response to *Yersinia pestis* infection. J Biol Chem 285, 10832–10840.

Bray, N.L., Pimentel, H., Melsted, P., and Pachter, L. (2016). Near-optimal probabilistic RNA-seq quantification. Nat Biotechnol 34, 525–527.

Brennan, J.J., and Gilmore, T.D. (2018). Evolutionary Origins of Toll-like Receptor Signaling. Mol Biol Evol 35, 1576–1587.

Brennan, J.J., Messerschmidt, J.L., Williams, L.M., Matthews, B.J., Reynoso, M., and Gilmore, T.D. (2017). Sea anemone model has a single Toll-like receptor that can function in pathogen detection, NF-κB signal transduction, and development. Proc Natl Acad Sci U S A 114, E10122–E10131.

Brenner, S. (1974). The genetics of *Caenorhabditis elegans*. Genetics 77, 71–94.

Cao, X., and Aballay, A. (2016). Neural Inhibition of Dopaminergic Signaling Enhances Immunity in a Cell-Non-autonomous Manner. Curr Biol 26, 2329–2334.

Cao, X., Kajino-Sakamoto, R., Doss, A., and Aballay, A. (2017). Distinct Roles of Sensory Neurons in Mediating Pathogen Avoidance and Neuropeptide-Dependent Immune Regulation. Cell Rep 21, 1442–1451.

Cezairliyan, B., Vinayavekhin, N., Grenfell-Lee, D., Yuen, G.J., Saghatelian, A., and Ausubel, F.M. (2013). Identification of *Pseudomonas aeruginosa* phenazines that kill *Caenorhabditis elegans*. PLoS Pathog 9, e1003101.

Chandra, V., Huang, P., Potluri, N., Wu, D., Kim, Y., and Rastinejad, F. (2013). Multidomain integration in the structure of the HNF-4α nuclear receptor complex. Nature 495, 394–398.

Cheesman, H.K., Feinbaum, R.L., Thekkiniath, J., Dowen, R.H., Conery, A.L., and Pukkila-Worley, R. (2016). Aberrant Activation of p38 MAP Kinase-Dependent Innate Immune Responses Is Toxic to *Caenorhabditis elegans*. G3 (Bethesda) 6, 541–549.

Choi, K.H., Kumar, A., and Schweizer, H.P. (2006). A 10-min method for preparation of highly electrocompetent *Pseudomonas aeruginosa* cells: application for DNA fragment transfer between chromosomes and plasmid transformation. J Microbiol Methods 64, 391–397.

Conte, D., Jr., MacNeil, L.T., Walhout, A.J.M., and Mello, C.C. (2015). RNA Interference in *Caenorhabditis elegans*. Curr Protoc Mol Biol 109, 26 23 21–26 23 30.

Deng, P., Uma Naresh, N., Du, Y., Lamech, L.T., Yu, J., Zhu, L.J., Pukkila-Worley, R., and Haynes, C.M. (2019). Mitochondrial UPR repression during *Pseudomonas aeruginosa* infection requires the bZIP protein ZIP-3. Proc Natl Acad Sci U S A 116, 6146–6151.

Dietrich, L.E., Price-Whelan, A., Petersen, A., Whiteley, M., and Newman, D.K. (2006). The phenazine pyocyanin is a terminal signalling factor in the quorum sensing network of *Pseudomonas aeruginosa*. Mol Microbiol 61, 1308–1321.

Dokshin, G.A., Ghanta, K.S., Piscopo, K.M., and Mello, C.C. (2018). Robust Genome Editing with Short Single-Stranded and Long, Partially Single-Stranded DNA Donors in *Caenorhabditis elegans*. Genetics 210, 781–787.

Dunbar, T.L., Yan, Z., Balla, K.M., Smelkinson, M.G., and Troemel, E.R. (2012). *C. elegans* detects pathogen-induced translational inhibition to activate immune signaling. Cell Host Microbe 11, 375–386.

Evans, R., O’Neill, M., Pritzel, A., Antropova, N., Senior, A., Green, T., Žídek, A., Bates, R., Blackwell, S., Yim, J., et al. (2022). Protein complex prediction with AlphaFold-Multimer. bioRxiv, 2021.2010.2004.463034.

Filipowicz, A., Lalsiamthara, J., and Aballay, A. (2021). TRPM channels mediate learned pathogen avoidance following intestinal distention. eLife 10:e65935.

Fire, A., Xu, S., Montgomery, M.K., Kostas, S.A., Driver, S.E., and Mello, C.C. (1998). Potent and specific genetic interference by double-stranded RNA in *Caenorhabditis elegans*. Nature 391, 806–811.

Fitzgerald, K.A., and Kagan, J.C. (2020). Toll-like Receptors and the Control of Immunity. Cell 180, 1044–1066.

Fletcher, M., Tillman, E.J., Butty, V.L., Levine, S.S., and Kim, D.H. (2019). Global transcriptional regulation of innate immunity by ATF-7 in *C. elegans*. PLoS Genet 15, e1007830.

Folick, A., Oakley, H.D., Yu, Y., Armstrong, E.H., Kumari, M., Sanor, L., Moore, D.D., Ortlund, E.A., Zechner, R., and Wang, M.C. (2015). Aging. Lysosomal signaling molecules regulate longevity in *Caenorhabditis elegans*. Science 347, 83–86.

Foster, K.J., Cheesman, H.K., Liu, P., Peterson, N.D., Anderson, S.M., and Pukkila-Worley, R. (2020). Innate Immunity in the *C. elegans* Intestine Is Programmed by a Neuronal Regulator of AWC Olfactory Neuron Development. Cell Rep 31, 107478.

Friesner, R.A., Banks, J.L., Murphy, R.B., Halgren, T.A., Klicic, J.J., Mainz, D.T., Repasky, M.P., Knoll, E.H., Shelley, M., Perry, J.K., et al. (2004). Glide: a new approach for rapid, accurate docking and scoring. 1. Method and assessment of docking accuracy. J Med Chem 47, 1739–1749.

Gao, J.L., Kwan, A.H., Yammine, A., Zhou, X., Trewhella, J., Hugrass, B.M., Collins, D.A.T., Horne, J., Ye, P., Harty, D., et al. (2018). Structural properties of a haemophore facilitate targeted elimination of the pathogen *Porphyromonas gingivalis*. Nat Commun 9, 4097.

Garcia-Reyes, S., Cocotl-Yanez, M., Soto-Aceves, M.P., Gonzalez-Valdez, A., Servin-Gonzalez, L., and Soberon-Chavez, G. (2021). PqsR-independent quorum-sensing response of *Pseudomonas aeruginosa* ATCC 9027 outlier-strain reveals new insights on the *PqsE* effect on *RhlR* activity. Mol Microbiol 116, 1113–1123.

Gerstein, M.B., Lu, Z.J., Van Nostrand, E.L., Cheng, C., Arshinoff, B.I., Liu, T., Yip, K.Y., Robilotto, R., Rechtsteiner, A., Ikegami, K., et al. (2010). Integrative analysis of the *Caenorhabditis elegans* genome by the modENCODE project. Science 330, 1775–1787.

Ghanta, K.S., and Mello, C.C. (2020). Melting dsDNA Donor Molecules Greatly Improves Precision Genome Editing in *Caenorhabditis elegans*. Genetics 216, 643–650.

Halgren, T. (2007). New method for fast and accurate binding-site identification and analysis. Chem Biol Drug Des 69, 146–148.

Han, S.K., Lee, D., Lee, H., Kim, D., Son, H.G., Yang, J.S., Lee, S.V., and Kim, S. (2016). OASIS 2: online application for survival analysis 2 with features for the analysis of maximal lifespan and healthspan in aging research. Oncotarget 7, 56147–56152.

Harder, E., Damm, W., Maple, J., Wu, C., Reboul, M., Xiang, J.Y., Wang, L., Lupyan, D., Dahlgren, M.K., Knight, J.L., et al. (2016). OPLS3: A Force Field Providing Broad Coverage of Drug-like Small Molecules and Proteins. J Chem Theory Comput 12, 281–296.

Haynes, C.M., Yang, Y., Blais, S.P., Neubert, T.A., and Ron, D. (2010). The matrix peptide exporter HAF-1 signals a mitochondrial UPR by activating the transcription factor ZC376.7 in *C. elegans*. Mol Cell 37, 529–540.

Higgins, D.P., Weisman, C.M., Lui, D.S., D’Agostino, F.A., and Walker, A.K. (2022). Defining characteristics and conservation of poorly annotated genes in *Caenorhabditis elegans* using WormCat 2.0. Genetics.

Holdorf, A.D., Higgins, D.P., Hart, A.C., Boag, P.R., Pazour, G.J., Walhout, A.J.M., and Walker, A.K. (2020). WormCat: An Online Tool for Annotation and Visualization of *Caenorhabditis elegans* Genome-Scale Data. Genetics 214, 279–294.

Irazoqui, J.E., Urbach, J.M., and Ausubel, F.M. (2010). Evolution of host innate defence: insights from *Caenorhabditis elegans* and primitive invertebrates. Nat Rev Immunol 10, 47–58.

Janeway, C.A., Jr. (1989). Approaching the asymptote? Evolution and revolution in immunology. Cold Spring Harb Symp Quant Biol 54 *Pt* *1*, 1–13.

Jorgensen, W.L., Maxwell, D.S., and Tirado-Rives, J. (1996). Development and testing of the OPLS all-atom force field on conformational energetics and properties of organic liquids. Journal of the American Chemical Society 118, 11225–11236.

Labun, K., Montague, T.G., Krause, M., Torres Cleuren, Y.N., Tjeldnes, H., and Valen, E. (2019). CHOPCHOP v3: expanding the CRISPR web toolbox beyond genome editing. Nucleic Acids Res 47, W171–W174.

Lee, D.G., Urbach, J.M., Wu, G., Liberati, N.T., Feinbaum, R.L., Miyata, S., Diggins, L.T., He, J., Saucier, M., Deziel, E., et al. (2006). Genomic analysis reveals that *Pseudomonas aeruginosa* virulence is combinatorial. Genome Biol 7, R90.

Liberati, N.T., Urbach, J.M., Miyata, S., Lee, D.G., Drenkard, E., Wu, G., Villanueva, J., Wei, T., and Ausubel, F.M. (2006). An ordered, nonredundant library of *Pseudomonas aeruginosa* strain PA14 transposon insertion mutants. Proc Natl Acad Sci U S A 103, 2833–2838.

Lin, C.J., and Wang, M.C. (2017). Microbial metabolites regulate host lipid metabolism through NR5A-Hedgehog signalling. Nat Cell Biol 19, 550–557.

Magner, D.B., and Antebi, A. (2008). *Caenorhabditis elegans* nuclear receptors: insights into life traits. Trends Endocrinol Metab 19, 153–160.

Mahajan-Miklos, S., Tan, M.W., Rahme, L.G., and Ausubel, F.M. (1999). Molecular mechanisms of bacterial virulence elucidated using a *Pseudomonas aeruginosa*-*Caenorhabditis elegans* pathogenesis model. Cell 96, 47–56.

Martinez Molina, D., Jafari, R., Ignatushchenko, M., Seki, T., Larsson, E.A., Dan, C., Sreekumar, L., Cao, Y., and Nordlund, P. (2013). Monitoring drug target engagement in cells and tissues using the cellular thermal shift assay. Science 341, 84–87.

Mavrodi, D.V., Blankenfeldt, W., and Thomashow, L.S. (2006). Phenazine compounds in fluorescent Pseudomonas spp. biosynthesis and regulation. Annu Rev Phytopathol 44, 417–445.

Mavrodi, D.V., Bonsall, R.F., Delaney, S.M., Soule, M.J., Phillips, G., and Thomashow, L.S. (2001). Functional analysis of genes for biosynthesis of pyocyanin and phenazine-1-carboxamide from *Pseudomonas aeruginosa* PAO1. J Bacteriol 183, 6454–6465.

McEwan, D.L., Kirienko, N.V., and Ausubel, F.M. (2012). Host translational inhibition by *Pseudomonas aeruginosa* Exotoxin A Triggers an immune response in *Caenorhabditis elegans*. Cell Host Microbe 11, 364–374.

Meisel, J.D., Panda, O., Mahanti, P., Schroeder, F.C., and Kim, D.H. (2014). Chemosensation of bacterial secondary metabolites modulates neuroendocrine signaling and behavior of *C. elegans*. Cell 159, 267–280.

Melo, J.A., and Ruvkun, G. (2012). Inactivation of conserved *C. elegans* genes engages pathogen-and xenobiotic-associated defenses. Cell 149, 452–466.

Motola, D.L., Cummins, C.L., Rottiers, V., Sharma, K.K., Li, T., Li, Y., Suino-Powell, K., Xu, H.E., Auchus, R.J., Antebi, A., et al. (2006). Identification of ligands for DAF-12 that govern dauer formation and reproduction in *C. elegans*. Cell 124, 1209–1223.

Moura-Alves, P., Fae, K., Houthuys, E., Dorhoi, A., Kreuchwig, A., Furkert, J., Barison, N., Diehl, A., Munder, A., Constant, P., et al. (2014). AhR sensing of bacterial pigments regulates antibacterial defence. Nature 512, 387–392.

Moura-Alves, P., Puyskens, A., Stinn, A., Klemm, M., Guhlich-Bornhof, U., Dorhoi, A., Furkert, J., Kreuchwig, A., Protze, J., Lozza, L., et al. (2019). Host monitoring of quorum sensing during *Pseudomonas aeruginosa* infection. Science 366.

Mukherjee, S., Moustafa, D., Smith, C.D., Goldberg, J.B., and Bassler, B.L. (2017). The RhlR quorum-sensing receptor controls *Pseudomonas aeruginosa* pathogenesis and biofilm development independently of its canonical homoserine lactone autoinducer. PLoS Pathog 13, e1006504.

Mukherjee, S., Moustafa, D.A., Stergioula, V., Smith, C.D., Goldberg, J.B., and Bassler, B.L. (2018). The PqsE and RhlR proteins are an autoinducer synthase-receptor pair that control virulence and biofilm development in *Pseudomonas aeruginosa*. Proc Natl Acad Sci U S A 115, E9411–E9418.

Papenfort, K., and Bassler, B.L. (2016). Quorum sensing signal-response systems in Gram-negative bacteria. Nat Rev Microbiol 14, 576–588.

Pellegrino, M.W., Nargund, A.M., Kirienko, N.V., Gillis, R., Fiorese, C.J., and Haynes, C.M. (2014). Mitochondrial UPR-regulated innate immunity provides resistance to pathogen infection. Nature 516, 414–417.

Peterson, N.D., Cheesman, H.K., Liu, P., Anderson, S.M., Foster, K.J., Chhaya, R., Perrat, P., Thekkiniath, J., Yang, Q., Haynes, C.M., et al. (2019). The nuclear hormone receptor NHR-86 controls anti-pathogen responses in *C. elegans*. PLoS Genet 15, e1007935.

Peterson, N.D., Icso, J.D., Salisbury, J.E., Rodriguez, T., Thompson, P.R., and Pukkila-Worley, R. (2022). Pathogen infection and cholesterol deficiency activate the *C. elegans* p38 immune pathway through a TIR-1/SARM1 phase transition. eLife 11:e74206.

Pfaffl, M.W. (2001). A new mathematical model for relative quantification in real-time RT-PCR. Nucleic Acids Res 29, e45.

Pimentel, H., Bray, N.L., Puente, S., Melsted, P., and Pachter, L. (2017). Differential analysis of RNA-seq incorporating quantification uncertainty. Nat Methods 14, 687–690.

Pukkila-Worley, R. (2016). Surveillance Immunity: An Emerging Paradigm of Innate Defense Activation in *Caenorhabditis elegans*. PLoS Pathog 12, e1005795.

Pukkila-Worley, R., Feinbaum, R., Kirienko, N.V., Larkins-Ford, J., Conery, A.L., and Ausubel, F.M. (2012). Stimulation of host immune defenses by a small molecule protects *C. elegans* from bacterial infection. PLoS Genet 8, e1002733.

Pukkila-Worley, R., Feinbaum, R.L., McEwan, D.L., Conery, A.L., and Ausubel, F.M. (2014). The evolutionarily conserved mediator subunit MDT-15/MED15 links protective innate immune responses and xenobiotic detoxification. PLoS Pathog 10, e1004143.

Qiu, D., Damron, F.H., Mima, T., Schweizer, H.P., and Yu, H.D. (2008). PBAD-based shuttle vectors for functional analysis of toxic and highly regulated genes in *Pseudomonas* and *Burkholderia* spp. and other bacteria. Appl Environ Microbiol 74, 7422–7426.

Rahme, L.G., Stevens, E.J., Wolfort, S.F., Shao, J., Tompkins, R.G., and Ausubel, F.M. (1995). Common virulence factors for bacterial pathogenicity in plants and animals. Science 268, 1899–1902.

Recinos, D.A., Sekedat, M.D., Hernandez, A., Cohen, T.S., Sakhtah, H., Prince, A.S., Price-Whelan, A., and Dietrich, L.E. (2012). Redundant phenazine operons in *Pseudomonas aeruginosa* exhibit environment-dependent expression and differential roles in pathogenicity. Proc Natl Acad Sci U S A 109, 19420–19425.

Saunders, S.H., Tse, E.C.M., Yates, M.D., Otero, F.J., Trammell, S.A., Stemp, E.D.A., Barton, J.K., Tender, L.M., and Newman, D.K. (2020). Extracellular DNA Promotes Efficient Extracellular Electron Transfer by Pyocyanin in *Pseudomonas aeruginosa* Biofilms. Cell 182, 919–932 e919.

Schiessl, K.T., Hu, F., Jo, J., Nazia, S.Z., Wang, B., Price-Whelan, A., Min, W., and Dietrich, L.E.P. (2019). Phenazine production promotes antibiotic tolerance and metabolic heterogeneity in *Pseudomonas aeruginosa* biofilms. Nat Commun 10, 762.

Schindelin, J., Arganda-Carreras, I., Frise, E., Kaynig, V., Longair, M., Pietzsch, T., Preibisch, S., Rueden, C., Saalfeld, S., Schmid, B., et al. (2012). Fiji: an open-source platform for biological-image analysis. Nat Methods 9, 676–682.

Shapira, M., Hamlin, B.J., Rong, J., Chen, K., Ronen, M., and Tan, M.W. (2006). A conserved role for a GATA transcription factor in regulating epithelial innate immune responses. Proc Natl Acad Sci USA 103, 14086–14091.

Simanek, K.A., Taylor, I.R., Richael, E.K., Lasek-Nesselquist, E., Bassler, B.L., and Paczkowski, J.E. (2022). The PqsE-RhlR Interaction Regulates RhlR DNA Binding to Control Virulence Factor Production in *Pseudomonas aeruginosa*. Microbiol Spectr 10, e0210821.

Singh, J., and Aballay, A. (2019a). Intestinal infection regulates behavior and learning via neuroendocrine signaling. eLife 8:e50033.

Singh, J., and Aballay, A. (2019b). Microbial Colonization Activates an Immune Fight-and-Flight Response via Neuroendocrine Signaling. Dev Cell 49, 89–99 e84.

Sluder, A.E., and Maina, C.V. (2001). Nuclear receptors in nematodes: themes and variations. Trends Genet 17, 206–213.

Sluder, A.E., Mathews, S.W., Hough, D., Yin, V.P., and Maina, C.V. (1999). The nuclear receptor superfamily has undergone extensive proliferation and diversification in nematodes. Genome Res 9, 103–120.

Styer, K.L., Singh, V., Macosko, E., Steele, S.E., Bargmann, C.I., and Aballay, A. (2008). Innate immunity in *Caenorhabditis elegans* is regulated by neurons expressing NPR-1/GPCR. Science 322, 460–464.

Subramanian, A., Tamayo, P., Mootha, V.K., Mukherjee, S., Ebert, B.L., Gillette, M.A., Paulovich, A., Pomeroy, S.L., Golub, T.R., Lander, E.S., et al. (2005). Gene set enrichment analysis: a knowledge-based approach for interpreting genome-wide expression profiles. Proc Natl Acad Sci U S A 102, 15545–15550.

Sural, S., and Hobert, O. (2021). Nematode nuclear receptors as integrators of sensory information. Curr Biol 31, 4361–4366 e4362.

Taubert, S., Ward, J.D., and Yamamoto, K.R. (2011). Nuclear hormone receptors in nematodes: evolution and function. Mol Cell Endocrinol 334, 49–55.

Timmons, L., Court, D.L., and Fire, A. (2001). Ingestion of bacterially expressed dsRNAs can produce specific and potent genetic interference in *Caenorhabditis elegans*. Gene 263, 103–112.

Troemel, E.R., Chu, S.W., Reinke, V., Lee, S.S., Ausubel, F.M., and Kim, D.H. (2006). p38 MAPK regulates expression of immune response genes and contributes to longevity in *C. elegans*. PLoS Genet 2, e183.

Warnhoff, K., Roh, H.C., Kocsisova, Z., Tan, C.H., Morrison, A., Croswell, D., Schneider, D.L., and Kornfeld, K. (2017). The Nuclear Receptor HIZR-1 Uses Zinc as a Ligand to Mediate Homeostasis in Response to High Zinc. PLoS Biol 15, e2000094.

Watson, E., and Walhout, A.J. (2014). *Caenorhabditis elegans* metabolic gene regulatory networks govern the cellular economy. Trends Endocrinol Metab 25, 502–508.

Williams, P., and Camara, M. (2009). Quorum sensing and environmental adaptation in *Pseudomonas aeruginosa*: a tale of regulatory networks and multifunctional signal molecules. Curr Opin Microbiol 12, 182–191.

Wu, Z., Isik, M., Moroz, N., Steinbaugh, M.J., Zhang, P., and Blackwell, T.K. (2019). Dietary Restriction Extends Lifespan through Metabolic Regulation of Innate Immunity. Cell Metab 29, 1192–1205 e1198.

Yammine, A., Gao, J., and Kwan, A.H. (2019). Tryptophan Fluorescence Quenching Assays for Measuring Protein-ligand Binding Affinities: Principles and a Practical Guide. Bio Protoc 9, e3253.

Yoneda, T., Benedetti, C., Urano, F., Clark, S.G., Harding, H.P., and Ron, D. (2004). Compartment-specific perturbation of protein handling activates genes encoding mitochondrial chaperones. J Cell Sci 117, 4055–4066.

Yunus, A.A., and Lima, C.D. (2009). Purification of SUMO conjugating enzymes and kinetic analysis of substrate conjugation. Methods Mol Biol 497, 167–186.

Zhang, L., Ward, J.D., Cheng, Z., and Dernburg, A.F. (2015). The auxin-inducible degradation (AID) system enables versatile conditional protein depletion in *C. elegans*. Development 142, 4374–4384.

Zugasti, O., Bose, N., Squiban, B., Belougne, J., Kurz, C.L., Schroeder, F.C., Pujol, N., and Ewbank, J.J. (2014). Activation of a G protein-coupled receptor by its endogenous ligand triggers the innate immune response of *Caenorhabditis elegans*. Nat Immunol 15, 833–838.

